# Expanding salivary biomarker detection by creating a synthetic neuraminic acid sensor via chimeragenesis

**DOI:** 10.1101/2024.06.13.598939

**Authors:** Samuel J. Verzino, Sharona A. Priyev, Valeria A. Sánchez Estrada, Ali GholamianMogaddan, Gemma X. Crowley, Alexandra Rutkowski, Amelia C. Lam, Elizabeth S. Nazginov, Paola Kotemelo, Agustina Bacelo, Jack D. Flannery, Ksenya Gavrilov, Desiree T. Sukhram, Frank X. Vázquez, Javier F. Juárez

## Abstract

Accurate and timely diagnosis of oral squamous cell carcinoma (OSCC) is crucial in preventing its progression to advanced stages with poorer prognosis. As such, the construction of sensors capable of detecting previously established disease biomarkers for the early and non-invasive diagnosis of this and many other conditions has enormous therapeutic potential. In this work, we apply synthetic biology techniques for the development of a whole-cell biosensor (WCB) that leverages the physiology of engineered bacteria *in vivo* to promote the expression of an observable effector upon detection of a soluble molecule. To this end, we have constructed a bacterial strain expressing a novel chimeric transcription factor (Sphnx) for the detection of N-acetyl-D-neuraminic acid (Neu5Ac), a salivary biomolecule correlated with the onset of OSCC. This WCB serves as the proof-of-concept of a platform that can eventually be applied to clinical screening panels for a multitude of oral and systemic medical conditions whose biomarkers are present in saliva.

## INTRODUCTION

Biosensors are analytical devices in which biological elements convert a physicochemical stimulus into a detectable signal^1,2^. Among the many existing variants, whole-cell biosensors (WCBs) stand out due to their versatility and engineering potential^1^. These are detection systems in which a physiologically active cell sustains an array of biomolecular elements (receptors, transporters, enzymes, regulators, etc.) capable of signal recognition, integration, and production of an observable output^1^. Among the organisms that can serve as WCBs, there are bacterial species which either produce or limit the production of reporter proteins in the presence of a target analyte^3,4^. Microbial WCBs have demonstrated potential in fields such as environmental monitorization, food safety, and medicine^3,5^. Microorganisms can grow rapidly in relatively inexpensive media and produce all the necessary biological elements for analyte detection, providing a low-cost and portable alternative to other detection systems^6^. Unlike conventional sensors that may require large and complex instruments as well as laborious sample processing, WCBs can detect a bioavailable substance directly from the environment with minimal preparation, enabling their incorporation into *in situ* detection hardware that may revolutionize point-of-care diagnosis^6^. Multiplexing strategies, which enable detection of several signals simultaneously, combined with improved safety controls (i.e. biocontainment strategies) increase the opportunities for these biosensors to be deployed in both clinical and diagnostic settings^3,4^. Additionally, the robust and versatile nature of bacterial biosensors makes them suitable for use in environments where other technologies would not be able to operate, such as strenuous pH and osmotic conditions^3^. However, the use of WCBs is not exempt from certain drawbacks, such as slow reaction times, potentially impractical dynamic and detection ranges, and sensitivity and selectivity issues. However, each year an increasingly large corpus of publications proposes solutions to these problems, making biosensors more competitive and reliable^7^.

WCBs benefit from the synergies resulting from the combination of native systems, evolved during millions of years to detect the miscellany of molecules surrounding microbes, with the ever growing toolkit that synthetic biology provides (such as the creation of engineered genetic circuits^8–11)^. In WCBs based on transcription factors (TFs) the presence of a soluble analyte becomes associated with a conformational change of a protein, so that the binding state of the polypeptide to DNA gets altered, modifying the transcription profile of a regulated reporter^12^. These biosensors benefit the most from the use of monogenic transcriptional repressors^8,9^, control systems encoded by a single gene whose product prevents transcription in the absence of a small soluble inducer molecule. The catalog of characterized TFs capable of interacting with small soluble molecules is expanding, yet still limited, which represents one of the main obstacles in the field^13,14^. To expand the panoply of molecules that can be detected, we can employ synthetic biology, whose technologies have enabled us to better understand protein modularity such that we can better engage in regulator customization and *de novo* creation^15,16^. In this paper, we centered our efforts on the detection of a salivary biomarker that, to the best of our knowledge, was not the inducer / inhibitor of any characterized TF and thus required the creation of a synthetic regulator.

Saliva is a biological fluid resulting from the secretions of several glands (among them the parotid, submandibular and sublingual), crevicular fluid from the gingiva, cell residues, microbes and associated structures such as biofilms, plus secretions from the respiratory tract^17,18^. Human saliva contains a vast array of metabolites informative of oral and systemic health^19^, making its frequent or continuous sampling by portable and wearable equipment extremely coveted. Translation of such sampling technologies to dental practices may increase their role at the forefront of early disease prevention and diagnosis^20^. Electrochemical intraoral biodevices capable of detecting salivary biomarkers presently exist, yet they are limited in the scope of molecules they detect and lack adaptability to sense new ones^21^. Biological detection systems based on engineered genetic circuits offer a plug-and-play potential, in which the substitution of one TF by another, plus its cognate operator, is often enough to repurpose a pre-existing sensor. Bioengineering solutions enabling point-of-care diagnostics will greatly benefit patients by improving at-home health monitorization^22^. This approach is especially relevant due to the widespread acceptance and use of consumer technology within society, which has normalized wearable biosensors connected to a phone or a smartwatch as well as at-home sample collection (e.g. for ancestry information)^23^. Diagnostic screenings that do not require invasive procedures (i.e. a blood sample or a biopsy) are preferred by patients, which makes saliva a prime biological fluid candidate to be sampled for health and disease markers^24,25^. In this work we focused our attention on N-acetyl-D-neuraminic acid (neuraminic acid, Neu5Ac), the most common of the sialic acids originally isolated from salivary glands^26–28^. Neu5Ac fulfills key roles in human physiology^29^ as well as in commensal and pathogenic bacteria^26^. Moreover, variation in salivary sialylation might become a suitable early indicator of changes associated with malignant pathologies, given their correlation to periodontal disease (PD) and oral cancer^30^. The oral microbiome of patients with PD include bacteria such as *Tannerella forsythia*, which engage in glycan harvesting behavior: they cleave Neu5Ac molecules from glycoproteins to stealthily evade their host’s immune system^31^. Additionally, abnormally high concentrations of Neu5Ac have been associated with the onset of oral cancer^32–34:^ a group of neoplasms which affect any region in the oral cavity, pharynx, and salivary glands^35^. The term oral cancer is often used interchangeably with oral squamous cell carcinoma (OSCC), which accounts for more than 90% of oral malignant epithelial neoplasms^35^. As one of the most prevalent cancers in the world^36^, untreated OSCC has very poor prognosis and, as such, early detection is essential for successful treatment^37^. Previous evaluations of sialic acid levels in the saliva of healthy individuals compared to patients with oral cancer provided a guide for the range of salivary Neu5Ac that can be observed. The concentration range identified in the saliva of healthy adults was approximately 21.65 ± 5.71 mg/dl (0.52 - 0.88 mM), while confirmed oral pre-cancer individuals displayed 59.75 ± 7.29 mg/dl (1.7 - 2.2 mM). Histopathologically confirmed OSCC patients showed a wide range of concentrations, 204.85 ± 60.38 mg/dl (4.6 - 8.6 mM), always superior to the other two groups^38^. Additional studies have employed multivariate regression analysis comparing both free and protein-bound salivary sialic acid concentrations between OSCC patients and healthy individuals. While free sialic acid levels were found to be significantly elevated in patients diagnosed with oral cancer, differences in protein-bound sialic acid levels were not found to be statistically significant^39^. These observations are further evidence that total salivary sialic acid concentration can be utilized as a biomarker for OSCC. It is important to note that there is at least one study in which Neu5Ac concentrations in saliva were decreased in OSCC patients, having been hypothesized that a downregulation of the pathways modulating Neu5Ac production might have occurred in these subjects^40^. Beyond its connection to OSCC, it has also been proposed that salivary Neu5Ac could interfere with SARS-Cov-2 (COVID-19) infection, acting as a competitive inhibitor blocking the viral antigens recognized by the host receptors^41^. Each of these factors made quantification of salivary Neu5Ac a desirable target for detection by biosensors.

The absence of a native TF capable of detecting Neu5Ac motivated us to construct a synthetic monogenic repressor for its integration in a genetic circuit that would enable the strain bearing it to behave as a WCB. We favor these regulators due to their simplicity, compact nature, and ease of integration into synthetic circuits^8,9,42^. The process of constructing our custom Neu5Ac-sensing regulators is based on streamlined gene fusion / chimeragenesis methods developed by our laboratory to detect soluble molecules^15^ . Here, we describe a Neu5Ac TF-based WCB that employs engineered bacterial machinery to fulfill the functions of the three basic modules present in any detection system: a sensing module, a processing unit, and an actuator. The living WCB can intake a Neu5Ac signal by expressing a novel synthetic transcription factor, Sphnx, that bridges detection and processing (a common feature of bioreceptors^23^). A separate reporter plasmid completes the system providing a gene encoding a fluorescent protein (sfGFP, henceforth stylized as GFP) whose expression is regulated by the synthetic TF Sphnx so that detectable changes in fluorescence occur upon the detection of soluble Neu5Ac. Sphnx is a member of a new combinatorial library of TFs assembled via a simplified chimeragenesis process. Important clues on inter-domain communication of modular proteins have started to be gathered by comparing Sphnx to non-functional members of the TF collection. Protein tertiary structure modeling of functional and non-functional regulators has contributed to establish comparisons between them, helping to elucidate intricate and frequently difficult to predict dynamics that may condition the success or failure of any chimeric TF.

## RESULTS AND DISCUSSION

### Design and Construction of Synthetic Transcription Factors

The functionality of the WCBs that we built in this work hinged on the obtention of a TF capable of binding Neu5Ac with efficacy and precision. The Neu5Ac-induced monogenic transcriptional repressor we intended to assemble included three modules: a N-terminus (Nt) DNA Binding Domain (DBD), a linker sequence (LNK), and a C-terminus (Ct) Ligand Binding Domain (LBD). The DBD specifically recognizes an operator sequence in the promoter it regulates, driving the expression of the reporter gene (*sfgfp*). The LBD is tasked with the recognition of Neu5Ac, providing specificity to the TF. Finally, the LNK sequence acts as a bridge, providing the right amount of flexibility/rigidity necessary for the transfer of information between DBD and LBD: steric changes in the LBD associated with Neu5Ac binding get translated into steric changes in the DBD effecting a change of its oligomeric state that results in the unbinding of the TF from its operator and thus in a de-repression of transcription. By using a gene fusion / chimeragenesis approach we generated a library of repressors with a common Nt-DBD-LNK-LBD-Ct architecture.

Our selected DBD was sourced from the paradigm of transcriptional regulators: the *lac* repressor or LacI^43^. Though naturally induced by lactose/IPTG, we exclusively employ the DBD of LacI recognizing the operator *lacO* present in the *Plac* promoter (as defined in^15^). We decided to use DBD-LacI because the *lac* repressor originated as a modular protein with two domains, and has been previously used in the construction of several functional chimeras^44–46^. It is important to remark that the exclusive use of DBD-LacI, as opposed to the full protein, ensures that chimeric proteins carrying this domain will recognize their target operator, yet makes them incapable of responding to native inducers.

Candidate LBD selection was accomplished through bibliographical research. Since the LBD was the domain that recognized the molecule of interest for which we built the biosensing strain, it could be sourced from any whole protein or protein domain as opposed to being co-opted from a previously documented regulator. One of our main sources of the LBDs we employ in our chimeragenesis pipeline are periplasmic binding proteins (PBPs)^47^. These are standalone subunits of transporters that act as molecular shuttles that assist in the traffic of small solutes from the outer to the inner membrane in Gram negative bacteria^48^. PBPs share a common ancestor with the LBD domains of the LacI / GalR family of regulators^45^. To build the synthetic TF we required for this project, we needed to identify LBD candidates that strongly bound our soluble molecule of interest (Neu5Ac). PBPs SatA and SiaP were selected for their binding affinity and specificity for the transport of Neu5Ac. SatA is a product of the *satABCD* (*<u>s</u>*ialic *<u>a</u>*cid *t*ransport) operon that encodes an ATP Binding Cassette (ABC) transporter^49^ from *Haemophilus ducreyi*^50^. An alternative *satABCD* system present in *Corynebacterium glutamicum* has been demonstrated to be essential for its growth using Neu5Ac as sole carbon source^51^. SiaP is a product of the *siaPQM* operon, which encodes a Tripartite ATP-independent periplasmic (TRAP) transporter in *Haemophilus influenzae*^52,53^. SiaP was the first TRAP-associated PBP whose structure was elucidated (in *Vibrio cholerae*, *Pasteurella multocida* and *Fusobacterium nucleatum*^52,54^). Structural comparisons and thermodynamic studies of SatA and SiaP suggest that similar affinities for Neu5Ac are achieved in the two PBPs through distinct mechanisms: one enthalpically driven and the other entropically driven. SiaP, as well as other TRAP transporters, are hypothesized to command an enthalpically driven ligand binding strategy due to the abundance of charged residues (Arg, Lys, Asp, and Glu) at their binding pocket. SatA, alternatively, is hypothesized to bind its substrate in what is considered an entropic approach because of the numerous hydrophobic and polar residues (Tyr, Phe, Gly, Leu, Asn, and Ser) at its binding pocket^55^. The structure of *H. influenzae* SiaP in the presence of Neu5Ac (PDB 3B50) reveals the ligand bound in a deep cavity with its carboxyl group forming a salt bridge with a highly conserved Arg residue^52^. Initial studies of SiaP isolated from *H. influenzae* displayed a binding *Kd* of ≈120 nM when affinity for Neu5Ac was tested^53^. Subsequent studies established a value closer to *Kd* ≈58 nM^52^. Similar binding affinity of SatA complexes has been identified with a *Kd* ≈133 nM^55^. The collection of empirical data supporting the ability of SatA and SiaP to bind Neu5Ac made them ideal candidates to become the LBD of a chimeric TF responsive to Neu5Ac.

The final piece of the chimeric gene puzzle consisted on establishing a connection between DBD and LBD domains. This task was performed by elements called linker (LNK) regions that tend to be less structured than the domains they connect^56^. In order to maximize the chances of transmitting the information of Neu5Ac binding to the DBD, so that it can detach from *lacO* and enable transcription, we decided to connect DBD and LBD via two different strategies: i) a direct linkage without a polypeptide linker (condition identify as LNK1 for uniformity purposes), or ii), by one out of four linkers characterized by having different biophysical properties (identified as LNK2 to LNK5; *Supp. Table 1*). It is important to note that while LNK1 does not contain a linker *per se*, the LBDs we chose, by virtue of being PBPs, are proteins that would be exported to the periplasm of the cell under natural circumstances, and thus contain a native signal peptide at their N-terminus. This kind of peptide appears to be unstructured for the most part, barring a short alpha-helix region^57^. We decided to include the signal peptides of both SiaP and SatA due to existent examples of functional chimeras containing similar structures, such as the glucose-inducible SLCP_GL_^46^ and the benzoate-inducible CbnR-ABE44898-OD^15^ synthetic TFs. At this stage, we had identified all the elements we needed to start assembling our new chimeric TFs.

dsDNA fragments encoding the DBD and all LNK-LBD variants were obtained as detailed in the *Materials and Methods* section. To initiate the construction of a library containing all the desired chimeric TFs, we performed several scarless assembly reactions with all the domain-encoding fragments and linearized expression vector pUC19RBS, followed by transformation in NEB5-alpha competent cells. Resulting clones were isolated and individually sequenced. This approach was performed twice. Chimeric combinations that were not present in the libraries were re-attempted to be individually cloned five times with no success, at which point we proceeded to put pause to their construction and move the project forward, subcloning the TF collection we had generated up to that point in time into the low-copy vector pCKTRBS, what generated the set of plasmids we collectively identified as pCKT-*Chimera*. We hypothesized that chimeras unsuccessfully cloned at this stage may be potentially deleterious for the cells. It is important to note that the expression of any heterologous protein is *per se* taxing on bacterial metabolism, and that TFs might be especially burdensome^58,59^. Chimeric constructions including DBD-LacI and SiaP/SatA seem to provoke a certain toxicity to the cell, albeit to different degrees depending on the specific domain combination. Indirect evidence supporting this hypothesis was observed when constructions expressing the chimeric TFs were transformed into *E. coli* MC4100 to assess their behavior in M63 minimal media with glycerol as the sole carbon source. Under these resource-limited conditions, stressed bacteria have more difficulties in thriving, and MC4100 cells bearing pCKT-*Chimera* vectors displayed a very modest and erratic growth consistent with our assumptions (data not shown). Table 1 contains the chimeric TFs built, assayed, and presented in this study, as well as a simplified interpretation of their functionality when grown in rich media. **Screening of transcription factors**. A functional *in vivo* screening was implemented using GFP as reporter. Plasmid pHC_DYOLacI-R (*Supp. Table 2*) was transformed into candidate strains expressing the chimeric TFs from pCKTRBS vectors under the control of TetR-aTc. These two plasmids expressed in the same strain constituted a genetic circuit (*Figure 1a*) that enabled a controlled expression of the chimera as well as its controlled induction. This circuit behaves as a B-or-not-A universal two input gate^60^ (*Figure 1b*), where there are two inputs (A, aTc supplementation; B, Neu5Ac supplementation) and one output (detection of GFP fluorescence). Based on the addition of the inputs we can define three states in the genetic circuit (*Figure 1c*): *Basal* (aTc^-^/Neu5Ac^-^), no input added, GFP is expressed from its promoter in pHC_DYOLacI-R; *Repression* (aTc^+^/Neu5Ac^-^), the presence of aTc induces TetR which is incapable of remaining attached to *PtetO* and thus the chimera is expressed from its corresponding pCKT-*Chimera* vector repressing the expression of GFP; *De-repression* (aTc^+^/Neu5Ac^+^), the chimera is expressed but if it is functional it is induced by Neu5Ac, de-repressing the promoter driving the expression of GFP. By comparing the behavior of the strain expressing the aforementioned circuit under these conditions, we can rule out non-functional synthetic TFs. GFP-associated fluorescence data displayed in this publication is always expressed as relative fluorescence (Rel.Fluor = Fluorescence arbitrary units / OD600) which includes a correction of fluorescence by the optical density of the culture in order to account for growth disparities among different strains and conditions. In the case that the chimeric protein was not a functional repressor, relative GFP-associated fluorescence (Rel.fluor) could be detected in the *Repression* state (ΔRel.fluor_basal-repressed_ tending to 0). On the other hand, if the chimera was able to bind to *lacO* on the reporter promoter yet incapable to recognize Neu5Ac with its LBD, or to transmit a LBD to DBD conformational change resulting in its detachment from the operator, we would have obtained a functional super-repressor^61^ unable to respond to its inducer and, thus, not a valid tool to detect Neu5Ac (ΔRel.fluor_(de-repressed)-repressed_ tending to 0). *Figure 1d* illustrates the expected GFP production along a growth curve for a strain expressing the screening genetic circuit that included an ideal chimera (ΔRel.fluor_(de-repressed)-repressed_ >0). It is important to note that even though Neu5Ac is a monosaccharide, its supplementation to rich growth media did not significantly increase the OD600 of the assayed strains, at least up to the tested 10 mM concentration (*Supp.* Figure 1). It is also important to note that repression by LacI, or LacI-derived chimeras, is rarely absolute so that a certain level of leakage, and thus of background reporter gene expression, is always expected^46,62^.

**Figure 1.**
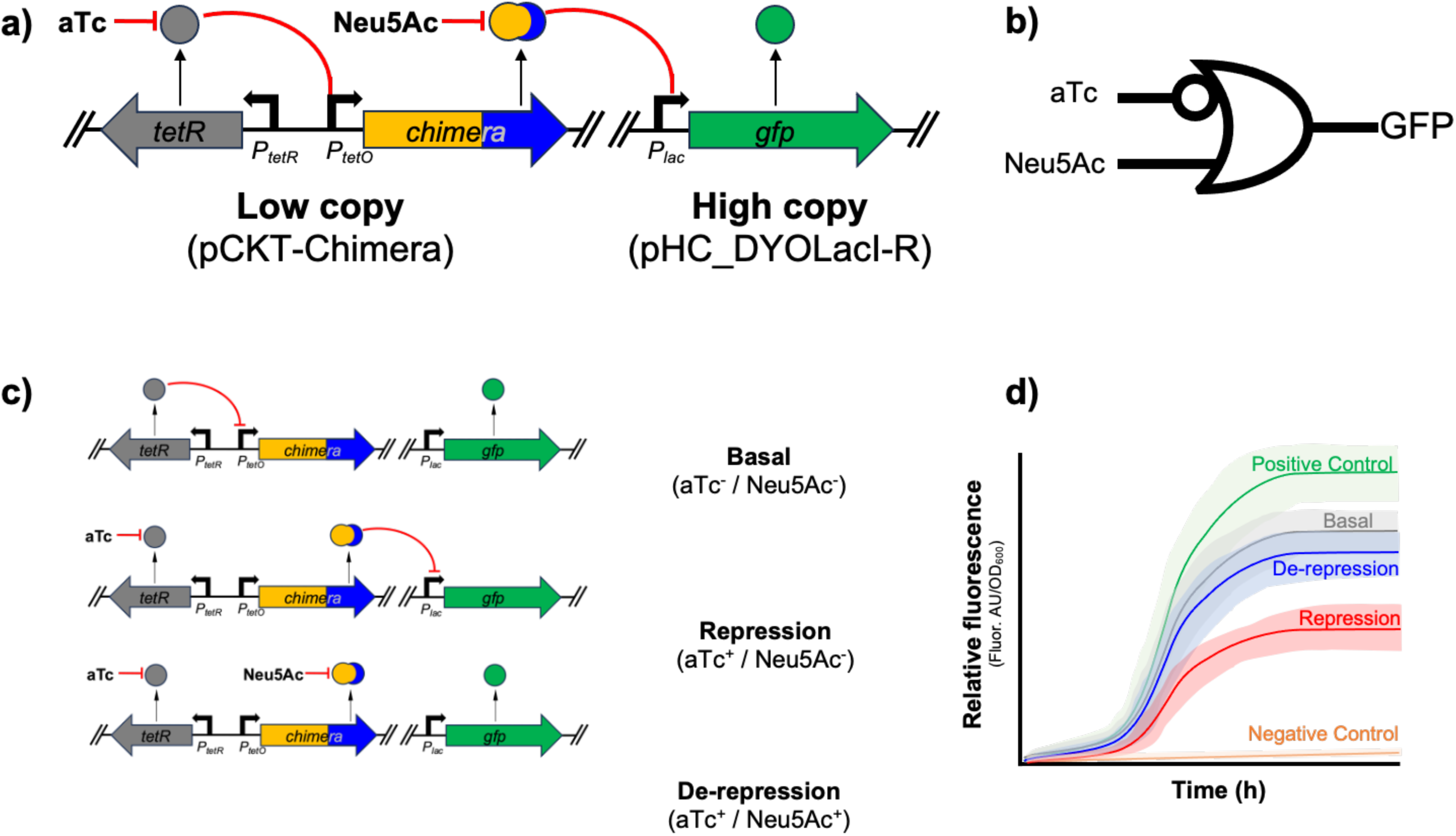
Screening strategy to find functional chimeric transcription factors. **a)** Biosensor genetic circuit for the *in vivo* testing of the functionality of cloned chimeric TFs. MG1655 (pCKT-*Chimera*, pHC_DYOLacI-R) cells express a chimeric TF under the control of TetR. The chimeric product can repress in *trans* the expression of a GFP reporter unless Neu5Ac is present. *tetR* / TetR (*grey arrow / circle*); chimeric TF gene / protein (*gold-blue arrow / circle*); *gfp* /GFP (*green arrow / circle*); anhydrotetracycline, aTc; neuraminic acid, Neu5Ac. **b)** Standard representation of the B-or-not-A universal two input gate that governs the expression of GFP in the circuit. **c)** States of the biosensor genetic circuit (top to bottom). *Basal*, the absence of aTc (aTc^-^) enables TeR to repress the expression of the chimeric repressor: regardless of the presence or absence of Neu5Ac GFP is expressed, and a fluorescent signal is identified. *Repressed*, the presence of aTc (aTc^+^) induces TetR which is unable to repress the promoter driving the transcription of the chimeric TF, which in turn represses the promoter driving the expression of *gfp* so that no fluorescent signal is detected. *De-repressed*, the presence of aTc (aTc^+^) ensures the production of the chimeric TF which, if functional, gets in turn induced by Neu5Ac, so that it cannot repress *gfp* and the GFP-associated fluorescence is detected again. c) Expected profile of fluorescent production over time by a bacterial culture expressing the biosensor genetic circuit and a functional chimeric TF. Relative fluorescence (GFP-associated fluorescence corrected by cell density measured as OD_600_). *Positive control* (*green line*), corresponds to the maximum GFP-associated fluorescence that can be observed, corresponding to a NEB5-alpha (pHC_DYOLacI-R) strain in which no repressor of the promoter driving GFP expression is present. *Negative control* (*orange line*), displays background fluorescence by a strain that does not carry *gfp*. *Basal* (*grey line*), *Repressed* (*red line*), *De-repressed* (*blue line*): GFP production associated to a MG1655 (pCKT-*Chimera*, pHC_DYOLacI-R) culture expressing a chimera in the basal, repressed, and de-repressed states, respectively.

**Table 1.** Chimeric transcription factors discussed in this work.

| Name |  | TF behaviour |  | Comments |
| --- | --- | --- | --- | --- |
| Common | Systematic | Repressor | De-repressible |  |
| Siren | LacI-LNK1-SiaP | Yes | Yes | Functional only in late stationary phase |
| Sphnx | LacI-LNK2-SiaP | Yes | Yes | Biosensor candidate |
| Kunst | LacI-LNK3-SiaP | Yes | No | - |
| Scorpio | LacI-LNK4-SiaP | No | No | - |
| Bastet | LacI-LNK5-SiaP | No | No | Repression in early hours of growth. |
| Lumos | LacI-LNK6-SiaP | No | No | Duplicated sequence acts as Linker. |
| Harpy | LacI-LNK1-SatA | Yes | Unreliable | Unstable strain |
| Haus | LacI-LNK2-SatA | No | No | - |

All the chimeric TFs included in Table 1 were screened for their ability to bind DNA and thus to repress the expression of the GFP reporter, as well as to be induced by Neu5Ac enabling production of the fluorescent protein. All members of the collection of MG1655 (pCKT-*Chimera*, pHC_DYOLacI-R) strains bearing the genetic circuit described in *Figure 1* were tested *in vivo* to assess their ability to behave as Neu5Ac biosensors (*Supp. Table 4*). Among the chimeric proteins expressed by this strain collection, we included the TF Lumos, which did not contain any of the expected LNK1-5 options but an unexpected 6 amino acid linker (LNK6) resulting from an abnormal assembly reaction in which the last 18 bases of *DBD-lacI* were duplicated (*Supp. Table 1*). Given the stable nature of the amino acid string, as well as our agnostic approach to which LNK sequence could be the best performing one, we decided to maintain this construction in our pool. TFs Kunst (LacI-LNK3-SiaP), Scorpio (LacI-LNK4-SiaP), Lumos (LacI-LNK6-SiaP), and Haus (LacI-LNK2-SatA) showed no repression abilities. These TFs seem to not be able to stably bind to their operator boxes in the reporter plasmid, potentially due to an inability to acquire their dimeric form. The strain bearing Haus also presented a diminished growth, which strongly suggests the potential toxicity of this transcription factor for the host cell.

Bastet (LacI-LNK5-SiaP) was intriguing due to the possibility that it was able to repress and de-repress for the first 10 hours of growth, yet its behavior requires a deeper study given the observed dispersion in the analyzed data. This observation suggests that the strain might be unstable due to the potential toxicity of the protein. Also of note, Harpy (LacI-LNK1-SatA) seemed to be able to repress at a certain level and potentially de-repress after the 18 hour mark. Another TF that presented this similar and unexpected induction profile but in more consistent basis was Siren (LacI-LNK1-SiaP), which displayed an intriguing function as a proper repressor inducible by Neu5Ac but only in the stationary phase of growth (*Supp.* Figure 2). In the future we intend to study this protein in depth, since its repression/de-repression profile is very attractive for a whole-cell biosensor setup in which a bacterial culture sustaining itself at stationary phase is continuously monitoring an input for an extended period of time. Prolonging the time WCBs are functional is a known challenge associated with these technologies, and engineering of genetic circuits incorporating regulators such as Siren, is one of the strategies that are being explored to account for this problem^3^. A candidate biosensor strain expressing Sphnx (LacI-LNK2-SiaP) exhibited the most suitable repression and induction profile compatible with that of a TF capable of repressing *Plac* while de-repressing it in a detectable manner upon binding to Neu5Ac. To summarize this initial evaluation of the functionality of our candidate TF-based WCBs, and judging by the difficulty to obtain them, their lack of functionality, and the reduced sample size, SatA-based fusions seem to be less stable and more prone to not be responsive to Neu5Ac. Further studies and the creation of expanded chimeric libraries, however, are required to confirm this hypothesis. At this injunction, we focused our efforts on better characterization of Sphnx, since it was the most promising TF we had assembled (estimated as the relative fluorescence difference under de-repressed and repressed conditions along the growth curve or (ΔRel.fluor_(de-repressed)-repressed)_ (*Supp.Table 4*).

In order to validate our preliminary observations, we systematically compared GFP production over time of MG1655 (pCKT-Sphnx, pHC_DYOLacI-R), MG-Sphnx for short, cells expressing Sphnx either in the presence or absence of Neu5Ac. *Figure 2a* displays the results of time course experiments over a period of 25 hours. GFP production induced by the presence of Neu5Ac was observed from the first hour of growth, with the Neu5Ac-induced cultures showing a tendency to display higher fluorescence than the repressed ones. At this point our attention was captured by Kunst as a model for a non-functional chimera in opposition to Sphnx. *Supp.* Figure 3 shows the behavior of Kunst, which exhibited a tendency to repress *Plac* at the very beginning of a 40 h time course, yet after the 3 h mark did not show any difference between induced and uninduced conditions. A preliminary evaluation of the structural differences between Sphnx and Kunst is included in its own section of *Results and Discussion*.

**Figure 2.**
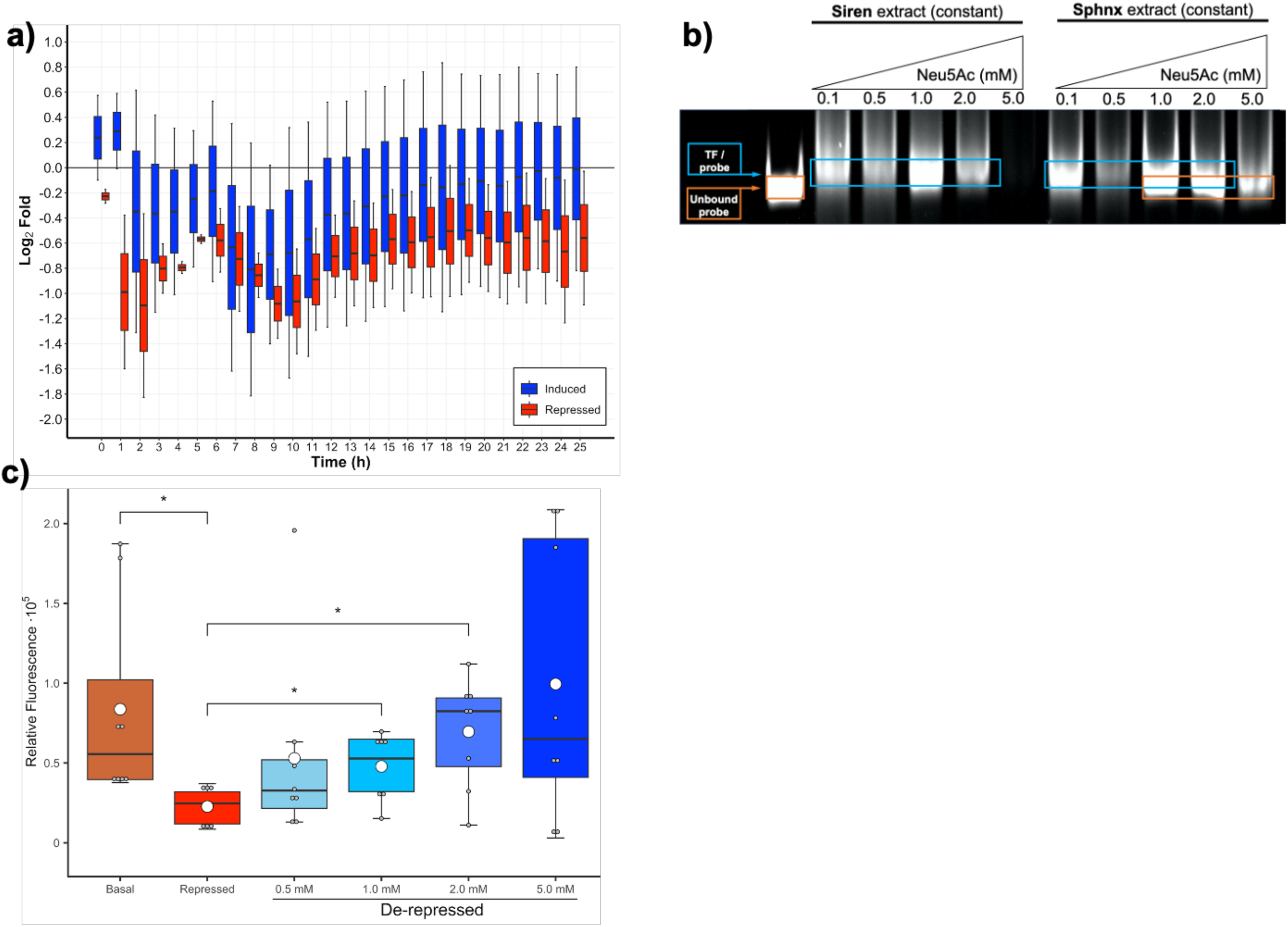
In vivo validation of the functionality of Sphnx (Lacl-LNK2-SiaP). a) Time course shcwng relative fluorescence of MG 1655 (pCKT-Sphnx. pHC_DYOLacf-R). MG-Sphnx. cells grown in a multiweB plate reader. Promoter activity is measured as bg_2_ fold of the relative fluorescence (GFP-associatcd fluorescence in arbitrary units I OD_ko_) of the strain growing in the do- repressed (aTc*. Neu5Ac*: blue boxplots) or repressed (aTc*. Neu5Ac; red boxplots) states compared to the basal expression of the reporter (aTc, Neu5Ac) Boxplots with whiskers represent data d-sperson of the average values of biological replicas (n = 8-10). It can be appreciated how in order to approach the basal activity (log? fold = 0). the addition of neuraminic aod is necessary when the chimera gets expressed by the supplementation of aTc Growth conditions and fluorescence assays performed as descnbed in Materials and Methods b) Sphnx and Siren protein interact with their cognate promoter P_w_. EMSA assays were performed as indicated under Materials and Methods. Interaction between NEB5-alpha (pCKT-Ch^nera) cell extracts expressing either Sphnx or Siren chlmenc TFs and a dsDNA probe (251 bp) including the P_w_ promoter. A fixed concentration of cell extract and DNA probe was exposed to growing concentrations of neuraminic acid (Neu5Ac). Lane numbers refer to the Nou5Ac concentration (mM) used for each reaction. The positions in the gel associated to unbound and bound probes are identified with orange aod Wue dashed rectangles. Respectively c) Relative fluorescence of MG-Sphnx cultures after induction with different concentrations of neuraminic acid and growth in a multrwell plate reader for 14 hours. Promoter activity is measured as Rel.fiuor, relative florescence (GFP-assooated fluorescence in arbitrary units / OD^_0_) of the strain growing in the basal (aTc. Neu5Ac; brown), repressed (aTc*. Neu5Ac: red) or de-repressed (aTc*. Neu5Ac*; blue shades) conditions. The concentrations of neuraminic acid (mM) added to the de­repressed conditions are specified under the corresponding plots. Boxplots with whiskers represent data dispersion of the average values of biological replicas (n = 8). The average value for every condition corresponds to a white orcle. Growth conditions, induction protocol and fluorescence assays performed as described in Melonals and Methods Statistically significant differences between selected conditions are marked with stars (p value < 0.06).

In an effort to validate the *in vivo* observations confirming Sphnx functionality, we analyzed the induction of *Plac* driving the expression of GFP in MG-Sphnx by qRT-PCR experiments (*Supp.* Figure 4) performed as indicated in *Materials and Methods*. The expression levels of *sfgfp* calculated as the ratio of the induced condition over the repressed condition were 1.58 ± 0.24 (*n* = 8, error expressed as SEM), statistically significant and consistent with an induction of the expression in the presence of Neu5Ac.

Further *in vitro* experiments were also performed to confirm the interaction between *Plac* and the DBD-LacI of Sphnx. Electrophoretic mobility shift assays (EMSA; *Materials and Methods*) showed that cell extracts from NEB5-alpha (pCKT-Sphnx) grown for 5 h were capable of retarding the migration of a *Plac-sfgfp* probe compared to control extracts that were not expressing Sphnx (data not shown). Neu5Ac behaved as the inducer of Sphnx, since binding of the protein to the probe was diminished in a concentration-dependent manner in the presence of the sialic acid (*Figure 2b*). On the other hand, cell extracts from NEB5-alpha (pCKT-Siren) grown for 5 h were able to bind the probe but not to unbind in the presence of Neu5Ac, confirming our *in vivo* observations. Previous publications report a similar behavior: DBDs from bacterial transcriptional repressors tend to retain their ability to bind to their operator sequences even when integrated in a new chimeric TF^63–65^, another testimony to the high modularity of these proteins. Our initial *in vitro* DNA-protein interaction studies require future validation with purified chimeras in order to calculate the kinetics of the Sphnx-*lacO* and Sphnx-Neu5Ac-*lacO* interactions, yet they represent a promising step forward towards characterizing the interaction between our synthetic TF and its cognate operator.

At this juncture of the project, both *in vivo* and *in vitro* experiments strongly suggested that Sphnx was a transcriptional repressor capable of interacting with *lacO* and being de-repressed by neuraminic acid. To better understand the dynamics of the Sphnx-Neu5Ac interaction we analyzed the production of GFP by MG-Sphnx cultures (*Materials and Methods*) when exposed to different concentrations of neuraminic acid around the average salivary concentration for an adult male (≈1 mM)^66^. The mean fold difference of GFP expression between the repressed and induced conditions was 2.25 ± 0.74 for cultures induced with 0.5 mM Neu5Ac, 2.54 ± 0.52 for cultures induced with 1.0 mM Neu5Ac, 3.81 ± 1.15 for cultures induced with 2.0 mM Neu5Ac, and 5.21 ± 2.00 for cultures induced with 5.0 mM Neu5Ac (*n* = 8, error expressed as SEM). These results were commensurable with those observed for modular chimeric TFs in the presence by their ligands: SLCPGL (LacI-GGBP-OD) induced by glucose^46^, CbnR-ABE44898-OD and LmrR-BzdB1_nSP induced by benzoate^15^. *Figure 2c* shows the levels of GFP fluorescence observed in basal and repressed cultures as well as in cultures induced by 0.5, 1.0, 2.0 and 5.0 mM neuraminic acid. These assays corroborated that relative fluorescence between basal (aTc^-^/Neu5Ac^-^) and repressed (aTc^+^/Neu5Ac^-^) conditions were significant. Cultures in which the genetic circuit used to test Sphnx were de-repressed (aTc^+^/Neu5Ac^+^) displayed statistically significant values compared to the repressed cultures when in the presence of 1.0 mM and 2.0 mM Neu5Ac. Induction by 0.5 and 5.0 mM was not significant. It is important to note that relative fluorescence values in the 5.0 mM-induced biological replicas were widely dispersed, suggesting that an overabundance of Neu5Ac might interfere with the stability of the host strain bearing both plasmids, producing some deleterious effect associated to the expression of the heterologous Sphnx and GFP proteins in the assayed conditions, which in practical terms means may entail the testing of a few sample dilutions when using these biosensors for out-of-the-lab applications.

In order to determine the specificity of Sphnx for Neu5Ac, we analyzed the production of GFP by MG-Sphnx cultures exposed to 1 mM Neu5Ac compared to cultures supplemented with 1mM of either of the following analogue compounds: N-acetyl-D-mannosamine (ManNAc), N-acetyl-D-glucosamine (GlcNAc), or N-glycolyl-neuraminic acid (Neu5Gc) (*Materials and Methods*). ManNAc and GlcNAc are aminated monosaccharides that are part of the biosynthetic pathway of Neu5Ac, constituting the backbone of the molecule^67–69^. Neu5Gc is a sialic acid that only differs from Neu5Ac in a single hydroxyl group and, interestingly, cannot be synthesized by humans, originating from diet and microbiota production^70–72^ Neu5Gc has been postulated as a cancer biomarker due to its proinflammatory effects^73,74^. Molecular docking simulations (*Figure 3a*) strongly suggested that both Neu5Ac and Neu5Gc could theoretically establish more stabilizing bonds with the amino acids present in the binding pocket of Sphnx-LBD, whereas ManNAc and GlcNAc would be able to enter said pocket yet their interaction with it would be extremely weak. These observations were corroborated *in vivo*. *Figure 3b* displays GFP-fluorescence levels by MG-Sphnx cultures exposed to the different Neu5Ac analogues. Induction by the different compounds is expressed as fold change Rel.fluor (*n* = 9-12, error expressed as SEM) of each induced culture (aTc^+^/ligand^+^) over a reference repressed condition (aTc^+^/ligand^-^). No statistically significant difference was found between repressed and ManAc^+^ or GlcNAc^+^ conditions, strongly suggesting that these compounds are not structurally similar enough for Sphnx to bind them. However, Neu5Gc^+^ cultures displayed a significant induction compared to the reference, presenting on average 62 ± 15% of the Neu5Ac^+^ induction. Upon further analysis there was no statistically significant difference between Neu5Ac^+^ and Neu5Gc^+^ datasets. These findings confirm that Neu5Gc is an inducer of Sphnx, which upon further review of the existing literature was coherent with previous observations that described how SiaP (Sphnx’s LBD) was able to bind Neu5Gc *in vitro* with μM affinity^55^

**Figure 3.**
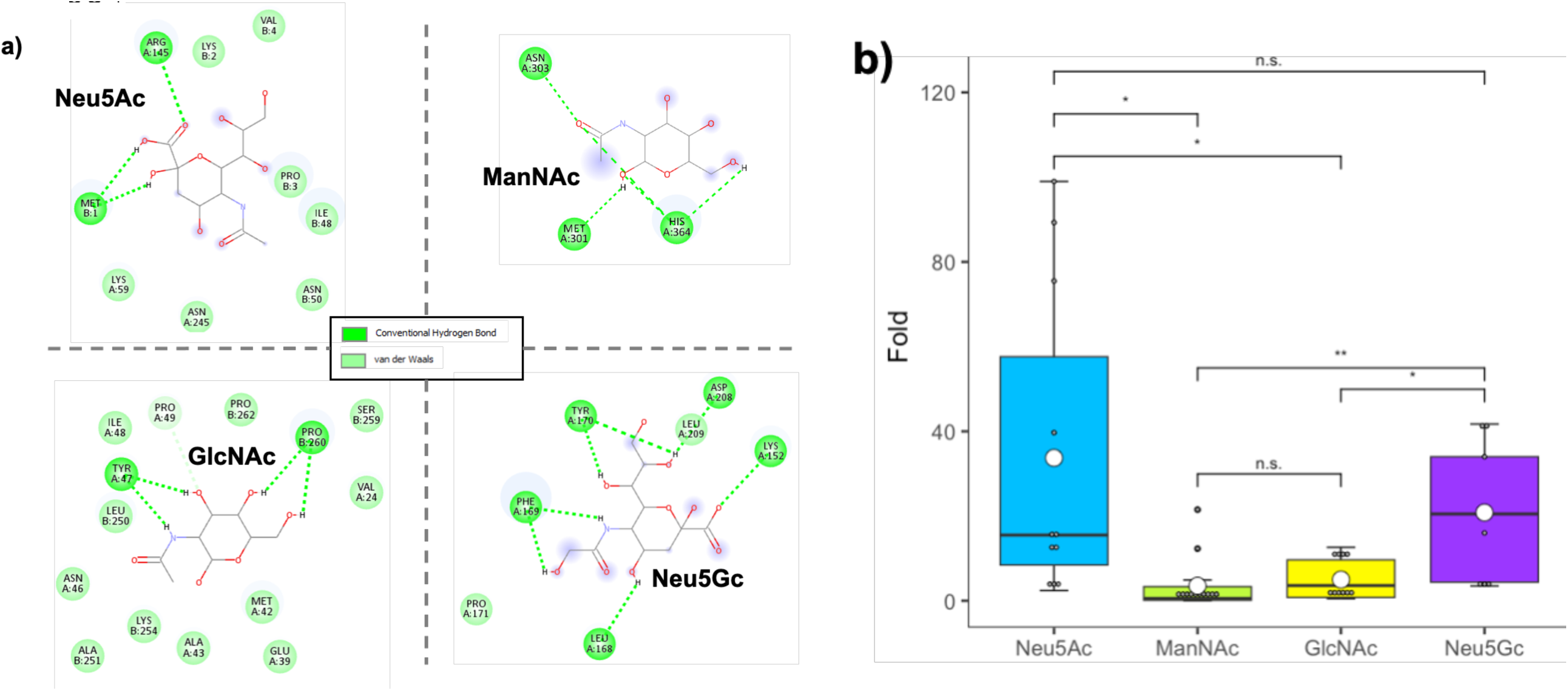
Interaction of Sphnx (LacI-LNK2-SiaP) with different ligands. Study of the interaction of Sphnx with N-acetyl-D-neuraminic acid eu5Ac), N-acetyl-D-mannosamine (ManNAc), N-acetyl-D-glucosamine (GlcNAc), and N-glycolyl-neuraminic acid (Neu5Gc). **a)** Molecular docking simulations for 3D models of Sphnx and ligand molecules displaying stabilizing van der Waals forces and hydrogen bonds between the binding pocket of the regulator and the ligands. Images generated in Discovery Studio (*Materials and Methods*). **b)** Relative fluorescence of MG1655 (pCKT-Sphnx, pHC_DYOLacI-R), MG-Sphnx, cultures after induction 1 mM of one of the following compounds: Neu5Ac (*blue*), ManNAc (*green*), GlcNAc (*yellow*), or Neu5Gc (*purple*) and growth in a shaker incubator with a subsequent transfer to a plate-reader. Promoter activity is measured as fold change of Rel.Fluor over a culture in the repressed condition (aTc^+^, ligand^-^). xplots with whiskers represent data dispersion of the average values of biological replicas (*n* = 9-12). The average value for every condition responds to a white circle. Growth conditions, induction protocol, and fluorescence assays performed as described in *Materials and thods*. Statistically significant differences between selected conditions are marked with stars (* *p* value < 0.05; ** *p* value < 0.01; n.s. non- nificant).

Our next objective was to test whether cultures of the WCB could detect Neu5Ac in contrived samples simulating human saliva (*Materials and Methods*). MG-Sphnx cultures mixed with artificial saliva supplemented with Neu5Ac for a final 1 mM concentration returned a Rel.Fluor signal 3.2 ± 0.9-fold (mean ± SEM, *n*=5) higher than samples of unsupplemented saliva. These data, even though preliminary, are very promising of the capabilities of WCBs, for diagnostic applications using clinical samples.

These collective results showing Sphnx induction are promising, yet they underline the need to refine chimeric TFs resulting from the assembly of pre-existing modules to optimize their induction range as well as gene dose and reporter genes used to study their behaviour *in vivo* and *in vitro*, as it has been previously proposed^14^. Newly created synthetic TFs tend to have limited transfer function and specificity, and thus to require further engineering for their field deployment in biosensors^75^. In this regard, we are optimistic about the future improvement of the chimeras we have assembled: the estimated *Kd* of SiaP and SatA for neuraminic acid are in the nM range^52,53,55^ (calculated *in vitro*), whereas in this work we present data in the mM range (calculated *in vivo*). This suggests that we have not reached the full potential of our LBDs and there is an opportunity to optimize Sphnx’s structure to better benefit from the high specificity and affinity of our LBD. It is also important to note that Sphnx is currently at the same point in evolutionary history when DBD-LacI was fused to the ancestral version of its current LBD, long before communication between DBD and LBD had been refined and perfected by natural selection^45^. A systematic exploration of promoter and RBS variants can provide a better solution to compensate for some of these deficiencies^14^. Regarding the use of alternative reporters that may shed more light on the induction profile of Sphnx, we acknowledge that the use of GFP has some inherent limitations in prokaryotes (e.g. widely varying maturation times dependent on strain and metabolic state^76^), which has motivated us to initiate the development of better reporting systems.

### Modeling of the 3D structure of chimeric transcription factors

Another element we are starting to explore is the importance of linkers for the functionality of modular proteins, an often-neglected design element^77^. Since the beginning of our journey characterizing the chimeric TFs, we found extremely intriguing the fact that Sphnx (LacI-LNK2-SiaP, Nt-DBD-EKEKEK-LBD- Ct) and Kunst (LacI-LNK3-SiaP, Nt-DBD-GSGSGS-LBD-Ct), which are nearly identical proteins (*Supp. Materials Appendix*), possessed opposed behaviours when interacting with neuraminic acid. The recent release of the AlphaFold 3 software^78^ has enabled us to predict the structures of multimeric chimeric proteins in the presence of their dsDNA operator, allowing us to model the proteins under conditions closer to those found in actual WCBs. Sphnx and Kunst were both modeled as dimers using AlphaFold 3, along with a DNA double helix containing the *lacO* (5’- ATTGTGAGCGGATAACAATT-3’ and its complementary sequence) recognized by the DBD- LacI they both carry. Our goal was to predict structures that could help explain why Sphnx binds Neu5Ac while Kunst does not, despite only differing by six amino acids. *Figure 4a* shows the predicted Kunst DBD (green) bound to the DNA double helix (gray) and the formation of a dimer between the PBP (cyan). The structure shows a mostly disordered linking region (orange), including the GSGSGS (LNK3) linker (violet). The predicted Sphnx model (*Figure 4b*), on the other hand, shows that the connecting region adopts a helical structure, including the EKEKEK (LNK2) linker, and appears to form contacts between both the DBD and PBP. Rotating the Kunst structure by 90° (*Figure 4c*) we can observe that the GSGSGS (linker does not interact with the protein or DNA. In contrast, rotating the Sphnx structure by 90° (*Figure 4d*) further shows that the linker forms contacts between the DBD and PBP domains, with interactions occurring across different monomer chains. Further analysis shows that the helical linker in Sphnx forms electrostatic salt bridges between itself and both the DBD and PBP domains. *Figure 4e* shows the acidic residues on the linker (Glu) forming contacts with basic residues (Lys) on the DBD and PBP domains. Specifically, the predicted structure suggests that the residues expected to form salt bridges include Glu67-Lys2 on the DBD and Glu69-Lys152 on the PBP. These salt bridges likely stabilize the helical linker, affecting the flexibility and stability of the polypeptide, an effect that has been previously observed in other proteins^79^. From these predicted structures, we hypothesize that these salt bridges stabilize helicity in the flexible linker in a way that positively affects the PBP function by interacting with the DBD and the PBP, explaining the ligand binding functionality of Sphnx compared to non-binding Kunst protein. Zooming out to see the larger structure (*Figure 4f*), aromatic residues near the linker salt bridges are visible. These residues are key for testing the new predicted structures and designing new chimeric proteins in the future. We intend to purify Sphnx and Kunst to perform fluorescence analysis of the protein-ligand interactions, including fluorescence quenching and fluorescence anisotropy studies. The construction of 3D models, using the latest version of AlphaFold, will enable us to explore which mutations might enhance ligand binding in a way that was previously unavailable, also allowing us to model conditions closer to the actual multimeric *in vivo* structures.

**Figure 4.**
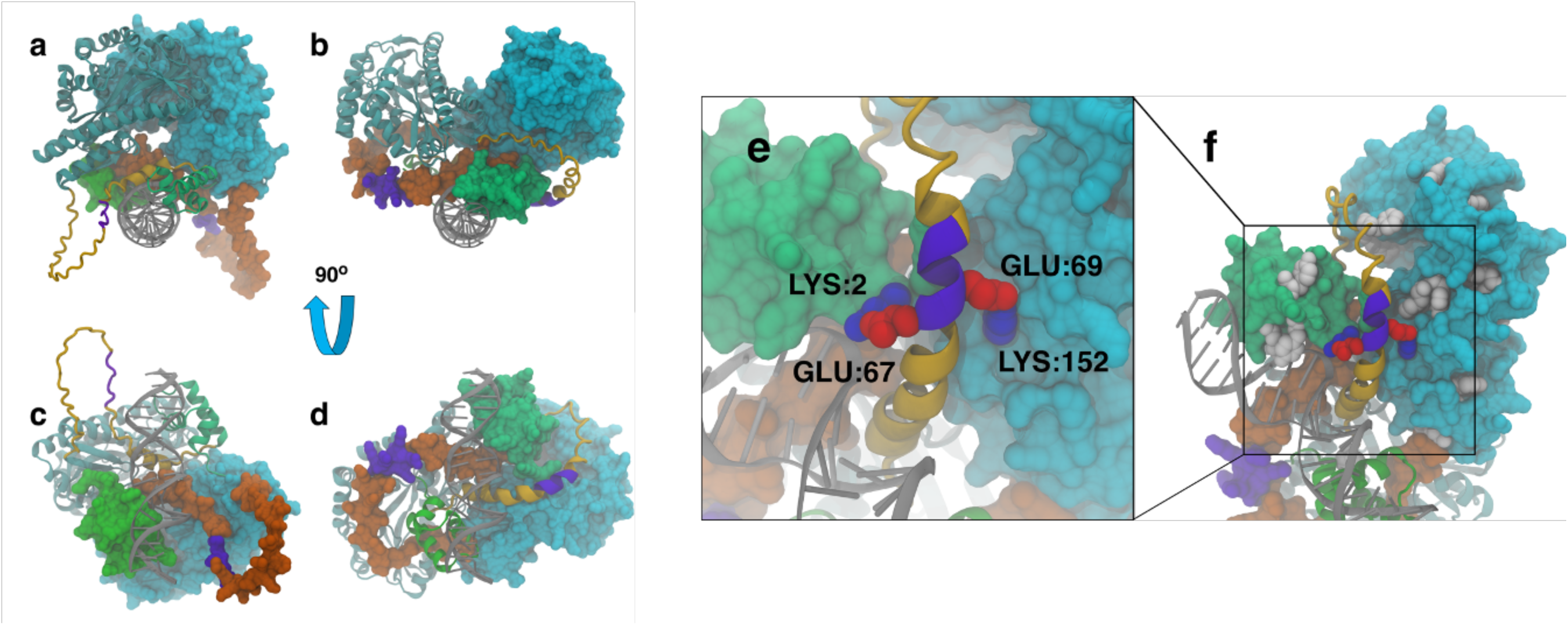
Modeled structures of chimeric protein dimers with double helix DNA obtained with Alphafold 3. For each structure, one monomer is shown using the New Cartoon representation and the other using the space-filling Surf representation. The DBD is shown in *green*, the PBP in *cyan*, and the connecting chain in *orange*, with the engineered linker in *violet*. The Kunst (**a**) and Sphnx (**b**) dimers are shown looking down the axis of the double helix, along with the rotated Kunst (**c**) and Sphnx (**d**) structures. The next panels contain the zoomed-in helical protein (**e**) linker (*violet*) in contact with the DBD (green) and PBP through the formation of salt bridges, with acidic residues shown in red and basic residues in blue. The zoomed-out protein (**f**) highlights the location of aromatic residues (*white*) that can be used to explore the validity of predicted structures using fluorescence anisotropy.

## CONCLUSIONS

This work presents MG1655 (pCKT-Sphnx, pHC_DYOLacI-R), MG-Sphnx, the first iteration of a transcription factor-based whole-cell biosensor strain capable of detecting the salivary biomarker N-acetyl-D-neuraminic acid and relaying its presence through the production of a fluorescent reporter, the traditional litmus test for a biosensor circuit^14^. The keystone of the genetic circuit enabling the detection of the soluble molecule is a new transcriptional repressor: Sphnx. The apparent absence of a native Neu5Ac-binding transcription factor led the authors to create a custom-made regulator via a straightforward synthetic biology approach that enables the generation of chimeras capable of repressing a known promoter and being de-repressed by a molecule of interest. Neuraminic acid is an attractive analyte to monitor since its enhanced levels in saliva are correlated with conditions that range from periodontal disease^66^ (oral microbes harvest sialic acid to evade our immune system^31^) to the onset of oral squamous cell carcinoma^32–34^. To the best of our knowledge, Sphnx is the first published transcriptional repressor capable of being induced by N-acetyl-D-neuraminic acid. Its ability to detect N-glycolyl-D-neuraminic as well makes MG-Sphnx a strong biosensor for the detection of sialic acid imbalances associated with multiple dysbiotic, inflammatory, and neoplasic conditions^74,80–82^.

This study supports the idea that the ability to construct custom-made TFs, affordably and on-demand, is instrumental in expanding the panoply of small soluble molecules that can be detected by biosensors in a field limited by the availability of sensing modules^14,45^. It is important to note that synthetic TFs such as Sphnx do not represent the closing of a chapter, but the origin of a new source of regulators whose performance can be improved via directed engineering of their ligand binding pockets^83,84^, an approach that can now benefit from high-throughput approaches^85^. It is widely acknowledged that biosensor specificity and biosensor-analyte reaction curves need to be optimized for the practical application of the WCB^75^. We have obtained promising results on the functionality of MG-Sphnx to detect N-acetyl-D-neuraminic in preparations of artificial saliva^86^, however, further experiments are needed to tune the parameters of the assay to detect a wide range of biomarker concentrations and, eventually, move on to test it with clinical salivary samples. The MG-Sphnx strain represents the initial step in the obtention of a functional WCB for the detection of neuraminic acid, a necessary launchpad to achieve the goal of eventual clinical deployment of the sensor.

The modular assembly of fusion genes such as Sphnx sheds light on the importance of inter-domain communication for multi-domain protein functionality^87^ and exemplifies how a reduced number of independent modules can be the playground of natural selection to synergically originate new functionalities in the cell^88^. The results presented in this work have challenged our expectations regarding chimeric transcription factors: while we expected most of them to be able to repress expression from their regulated promoter, we found several instances in which the chimeric product was unable to bind to its cognate operator boxes. Our current hypothesis is that different linkage alternatives condition the 3D structure of the multimodular protein to such an extent that the dimerization step necessary to bind the operator boxes cannot be achieved. We are looking forward to expanding these observations in the future. Our study presents the interesting contrast between the inducibility of Sphnx as opposed to Kunst, and particularly the unexpected role that a potential LBD-mediated stabilization of the linker may play in the de-repressibility of the TF dimer. *In vitro* assays using purified proteins, as well as further modeling simulation analysis will contribute to illuminate the complex dynamics necessary for modular proteins to fulfill their function. The eventual crystallization of these chimeras will provide the final clues on the matter.

In summary, this work presents a new biosensor for N-acetyl-D-neuraminic acid, a salivary biomarker relevant to oral health, which is based on a synthetic TF whose construction adds to our understanding on the role that different domains play in the functionality of modular proteins.

## MATERIALS AND METHODS

### Bacterial strains and growth conditions

Bacterial strains and plasmids used are listed in *Supp. Table 2*. *E. coli* strains were grown at 37°C in LB (Lysogenic Broth^89^) medium unless otherwise indicated. When required, *E. coli* cells were grown in M63 minimal medium^90^ using the necessary nutritional supplements and 30 mM glycerol (Sigma, St. Louis, MO) as a carbon source. Antibiotics were added at the following concentrations: 100 μg·ml^−1^ ampicillin/carbenicillin and 25 μg·ml^−1^ chloramphenicol (Sigma, St. Louis, MO). For protein expression from pCKTRBS- derived vectors, cultures were induced with 0.5 μg·ml^−1^ anhydrotetracycline (aTc) (Clontech, Mountain View, CA). When experimental conditions required it, cultures were supplemented with N-acetyl-D-neuraminic acid (neuraminic acid, Neu5Ac; Sigma, St. Louis, MO), N-acetyl-D-mannosamine (ManNAc; TCI Chemicals, Portland, OR), N-acetyl-D-glucosamine (GlcNAc; TCI Chemicals, Portland, OR), or N-glycolyl-neuraminic acid (Neu5Gc; AFB Bioscience-VWR, Radnor, PA) for a 1 mM concentration unless otherwise indicated.

Culture growth was routinely monitored in a BioPhotometer D30 (BioSpectrometer Basic Kinetic and Fluorescence; Eppendorf, Hamburg, Germany). When it was required, bacterial growth and GFP production were recorded in a Varioskan LUX Multimode Microplate Reader (Thermo Fisher, Waltham, MA). Cultures were pipetted on a 96-well plate (Flat bottom Polystyrene Black with clear bottom; Corning, Corning, NY) covered with a Breathe-Easy BEM-1 gas-permeable membrane (Diversified Biotech, Dedham, MA) and incubated in a plate reader, where OD600 and fluorescence in the range of GFP (excitation 485 nm, emission 528 nm) were either tracked along time courses (up to 40 h) or subjected to point measurements. NEB5-alpha (pHC_DYOLacI-R) and MG1655 were included in every plate as GFP-positive and GFP-negative reference strains, respectively. For standard time course experiments, overnight precultures of the strains were used to inoculate (starting OD600 0.025) fresh growth media (supplemented with aTc and Neu5Ac when indicated) in 96-well plates (180 μl total volume per well). For the single point measurements described in *Figure 2c* in which the MG-Sphnx is exposed to different concentrations of Neu5Ac, 5 ml overnight precultures of the strains (plus aTc and Neu5Ac when required) were used to inoculate 5 ml of fresh media supplemented in the same manner as the precultures. After 14 h of growth, 180 μl samples of the cultures were transferred to 96-well plates to enable OD600 and fluorescence readings. For the single point measurements described in *Figure 3b* in which the MG- Sphnx is exposed to 1mM Neu5Ac or 1mM of analogue compounds (ManNAc, GlcNAc, Neu5Gc), smaller 2 ml overnight precultures of the strain (plus aTc and Neu5Ac / ManNAc / GlcNAc / Neu5Gc when required) were used to innoculate 2 ml of fresh media supplemented as the precultures. After 20 h of growth, 180 μl samples of the cultures were transferred to 96-well plates to enable OD600 and fluorescence readings. For the experiments using artificial saliva a 5 ml overnight preculture of MG-Sphnx supplemented with aTc was used to inoculate (with 20 μl) a mixture of 750 μl of fresh media (plus aTc) and 250 μl of artificial saliva (artificial saliva with mucin, stabilized, Ref. 1700-0316; Pickering Laboratories, Mountain View, CA). The mix was incubated for 22 h at 37°C in an orbital shaker to ensure proper mixing and favor strain growth. This prolonged incubation seems to be key for the homogenization of the sample, since initial attempts to grow the mixture of culture and artificial saliva under the more gentle shaking conditions provided by the plate reader failed, presumably due to the viscosity of the saliva. Once the incubation was finished 180 μl samples of the cultures were transferred to 96-well plates to enable OD600 and fluorescence readings as described above for endpoint measurements. Incubation of the samples in the plate reader, after the endpoint measurement, with up to 48 h continuous OD600 and fluorescence monitorization did not improve the quality of the data.

Statistical analysis as well as data visualization were performed with R (https://www.r-project.org/) in the RStudio (RStudio Inc., Boston, MA) and Visual Studio Code environments (Microsoft Corp., Redmond, WA).

### Molecular biology techniques

Molecular biology techniques were performed following commonly used standard protocols and as per manufacturers’ instructions^91^. PCR reactions took place in a 6321 Mastercycler PRO Vapo Protect Thermal Cycler (Eppendorf, Hamburg, Germany). Routine separation of DNA and RNA in agarose gels was performed in an RunOne Electrophoresis system (EmbiTec, San Diego, CA). Agarose gels were imaged in an UVP UVSolo Touch (Analytic Jena, Jena, Germany). Plasmid DNA was purified with a Qiaprep Spin Miniprep Kit (Qiagen, Hilden, Germany). DNA fragments were purified with DNA Clean-up and Concentration Kit (Zymo research, Irvine, CA). The oligonucleotides employed for PCR amplification of the cloned fragments and other molecular biology techniques are summarized in *Supp. Table 3* and were supplied by IDT (Coralville, IA). All cloned inserts and DNA fragments were confirmed either via Sanger sequencing^92^ performed by Azenta Inc. (Burlington, MA), or Oxford-Nanopore whole-plasmid sequencing performed by Plasmidsaurus (Eugene, OR). Commercially available NEB5- alpha chemically competent cells (NEB, Ipswich, MA) were used for routine transformations. Alternatively, electrocompetent *E. coli* cells were generated and transformed immediately (Gene Pulser; Bio-Rad, Hercules, CA)^91^. Nucleotide sequence analyses were done at the National Center for Biotechnology Information server (www.ncbi.nlm.nih.gov) and Benchling Biology Software (www.benchling.com). Cloning was routinely performed via Gibson assembly^93^ unless otherwise indicated.

### Domain selection and cloning

DBD-LacI is located on *lacI* 5’ region, from the start codon to base 198, encoding a peptide of 66 amino acids that comprises the N-terminus of our chimeric proteins. This domain was obtained via PCR amplification from plasmid pCKTRBS-LacIwt^15^ using the oligonucleotide pair SV00001/SV00002. SatA from *H. ducreyi* (Uniprot Q7VL18) and SiaP from *H. influenzae* (Uniprot P44542) were ordered as synthetic dsDNA fragments (IDT, Coralville, IA). Several variants of the LBDs SatA and SiaP were amplified via PCR so that they would include 5’ overhangs including linkers (LNK2-5) or no linker whatsoever (LNK1). SatA was amplified with primers SV00030-SV00034 / SV00004 and SiaP with primers SV00015- SV00019 / JFJ0018.

Genes encoding the chimeric TFs were originally assembled in pUC19^94^. The vector was amplified via divergent PCR with primers JFJ00019/JFJ00020, which introduce a consensus RBS for *E. coli*, originating plasmid pUC19RBS. Tripartite Gibson assemblies including linear pUC19RBS, DBD- LacI, and LNK-LBD were performed and the resulting constructions transformed into NEB5-alpha competent cells. After a carbenicillin selection in solid media, a subset of the resulting clones was isolated and their plasmids (pUC19-*Chimera* family, *Supp. Table 2*) purified and sequenced to validate the presence of chimeric TFs. The resulting TF genes cloned into pUC19RBS were amplified using SV00024/SV00025 for SiaP-derived chimeras and SV00024/SV00026 for their SatA counterparts. These primers amplify our TF genes from the 5’ DBD-LacI, just downstream of the consensus RBS site, to the stop codon of the LBD. Low copy plasmid pCKTRBS^15^ was amplified via divergent PCR using primer pairs SV00022/SV00021 for SiaP-derived chimeras and SV00022/SV00023 for their SatA counterparts. These primers introduce 5’ and 3’ overhang sequences providing homology to our dsDNA chimeric gene fragments, and including in their 5’ side our consensus RBS site. Bipartite Gibson assemblies including linear pCKTRBS and the fusion genes encoding the chimeras were performed, and the resulting assembly products transformed into NEB5-alpha competent cells. After a carbenicillin selection in solid media, resulting clones were isolated and their plasmids sequenced to confirm the presence of chimera-encoding genes. Resulting pCKT-*Chimera* plasmids were purified from NEB5-alpha (pCKT- *Chimera*) strains and electroporated into MG1655 cells. After confirming the presence of the plasmid in the new host, MG1655 (pCKT-*Chimera*) were electroporated with the reporter plasmid pHC_DYOLacI-R. Validated MG1655 (pCKT-*Chimera*, pHC_DYOLacI-R) strains were used for *in vivo* and *in vitro* analysis of the functionality of the novel chimeric TFs when required. The sequences of all the oligonucleotides used in this work can be found in *Supp. Table 3*.

### Electrophoretic mobility shift assay (EMSA)

In order to obtain cell extracts enriched in the Sphnx/Siren proteins, NEB5-alpha (pCKT-Sphnx) and NEB5-alpha (pCKT-Siren) cells were grown on LB supplemented with aTc for 5 hours at 37°C in a shaker incubator. Cells were recovered by centrifugation at 4°C (15’ at 4,000 g) and then resuspended in 20 mM Tris-HCl (pH 7.0) to be sonicated at 70 amp for 30 seconds (30 cycles of 1 second; Branson Digital Sonifier, Danbury, CN). A subsequent centrifugation at 4°C (20’ at 20,000 g) was performed to eliminate cell debris from the sample. The recovered supernatant was then incubated with the DNA probe, a 251 bp dsDNA fragment amplified from plasmid pHC_DYOLacI-R with primers SV00028/SV00029, containing *Plac* (including *lacO*). Cell extract (5 μl) and DNA probe (5 μl) were incubated together with reaction buffer (20% glycerol, 100 mM KCl, 500 μg·ml^−1^ bovine serum albumin, 20mM Tris-HCl pH 7.5). The reaction mixture is incubated for 30’ at 30 °C. After incubation of the retardation mixtures, 2 µl of Gel loading dye without SDS (NEB, Ipswich, MA) were added to each reaction. Mixtures were subsequently fractionated by electrophoresis in precast 6% polyacrylamide gels (Novex TBE Gels; Invitrogen, Waltham, MA) buffered with 0.5X TBE (45 mM Tris borate, 1mM EDTA). Gels were run at constant voltage (100 V) for 90’ in a Mini Gel Tank (Invitrogen, Waltham, MA) and afterwards stained for 1 h in 50 ml of 0.5X TBE buffer supplemented with 5 µl of SYBR Gold dye (Invitrogen, Waltham, MA) while gently shaken. Post staining, gels were imaged in a UVP UVSolo Touch (Analytic Jena, Jena, Germany).

### qRT-PCR analysis

Quantitative Real Time Polymerase Chain Reaction (qRT-PCR) was used to assay the transcription of our reporter gene (*sfgfp*). MG1655 (pCKT-*Chimera*, pHCDYOLacI-R) strains were precultured overnight in of 3 ml LB supplemented with the corresponding selection antibiotics. On the next day, 100 µl of overnight preculture were used as inoculum for several cultures, 10 ml each, of fresh LB (plus selection antibiotics). Each one of the cultures was supplemented so that it would represent one of these three different conditions: *Basal* (aTc^-^, Neu5Ac^-^), *Repressed* (aTc^+^,Neu5Ac^-^), and *De-repressed* (aTc^+^, Neu5Ac^+^). Where experimental conditions required it, cultures were supplemented with 0.5 μg·ml^−1^ anhydrotetracycline (aTc) (Clontech, Mountain View, CA) and 1 mM neuraminic acid (Neu5Ac; Sigma, St. Louis, MO). For every independent qRT-PCR experiment, three biological replicates were created across each of these three conditions. These cultures were placed in a shaker incubator at 37°C for 5 hours and pelleted at 4°C and 4,000 g. Pellets were either frozen at -80°C for future use or immediately processed for total RNA extraction. Cells from each individual pellet were set for lysis by resuspension in 1 ml of IBI reagent, a phenol and guanidine isothiocyanate mixture (IBI Scientific, Dubuque, IA). 200 µl of chloroform (Fisher Scientific, Pittsburgh, PA) was added to the lysate and the mixture was vigorously shaken to homogeneity, followed by centrifugation (benchtop minifuge) at 4°C and 14,500 rpm for 15 minutes that led to a phase separation. 500 µl of the upper aqueous phase was mixed with an equal volume of 100% ethanol. At this point, total RNA was purified from DNA/RNA phase with Direct-zol RNA Miniprep Kit (Zymo Research, Irvine, CA) where the TRIzol reagent steps are replaced with the phenol and chloroform phase separation. Total RNA eluted with ddH2O was then retrotranscribed using qScript Flex cDNA Synthesis Kit (QuantaBio, Beverly, MA). A volume totaling 1 µg RNA was incubated with random hexamers and then supplemented with retrotranscriptase enzyme mix to synthesize cDNA. cDNA resulting from the last step was used as a template for the amplification of a 256 bp fragment with oligonucleotides SV00027A/SV00027 in a CFX96 Real-Time PCR System (Bio-Rad, Hercules, CA). Primers had been designed to amplify a region of pHC_DYOLacI comprising the beginning of *sfgfp*, so that when the tested chimera was being expressed it repressed the expression of GFP unless Neu5Ac was present. Cq values were analyzed as a measure of gene expression using the ΔΔCq (2^-ΔΔCt^) method^95^.

### Protein Structure Prediction using AlphaFold 3

The 3D structures of the target proteins, Sphnx (LacI-LNK2-SiaP, Nt-DBD-EKEKEK-LBD-Ct) and Kunst (LacI-LNK3-SiaP, Nt-DBD-GSGSGS-LBD-Ct), were predicted using AlphaFold 3^78^, developed by Google DeepMind (London, UK). The predictions were performed using the publicly available AlphaFold 3 web server (https://golgi.sandbox.google.com/). Both Sphnx and Kunst proteins were modeled as dimers, along with the DNA double helix containing *lacO* (5’-ATTGTGAGCGGATAACAATT- 3’ and its complementary sequence). Post-prediction, the structures were visualized and further analyzed using VMD^96^ to determine key protein-protein interactions.

### Molecular docking simulations

Sphnx (LacI-LNK2-SiaP) and Kunst 3D structures were used to forecast the primary binding patterns between the TF and a collection of putative ligands: Neu5Ac (PubChem CID 439197), ManNAc (PubChem CID 439281), GlcNAc (PubChem CID 439174), and Neu5Gc (PubChem CID 440001). First, water molecules and heteroatoms were removed from the PDB files for Sphnx and Kunst 3D structure. Using AutoDock4^94^, Kollman charges were supplemented to the 3D structure of both our ligands (see PubChem reference above) and proteins, with polar hydrogen charges being also added to the transcription factor structures. Once ligands and receptors were prepared, a grid box delineating the region of the whole protein where the docking would take place was established. Employing the molecular docking simulation protocols provided by the AutoDock 4 suite, binding affinities were assessed. Post-docking, visualization software (Pymol, Schrödinger Inc., New York, NY; Discovery Studio, Biovia, San Diego, CA) were used to identify interactions between Sphnx / Kunst and the aforementioned ligands.

## Acknowledgments and Funding

This material is based upon work supported by St. John’s University (SJU) via J.F.J.’s starting package as well as St. John’s SEED grants (2020-2021, 2021-2022, 2024-2025). Analysis and visualization of predicted structures was performed by F.X.V and A.B. on hardware resources supported by the National Institutes of Health under award number NIH SC2GM131992. This work is solely the responsibility of the authors and does not necessarily represent the official views of St. John’s University. We want to acknowledge L. Bryan, D. Howarth, T. Mustacchio, M. Ruggiu, and W. Munyao (SJU) for their support and time sharing equipment, reagents, and protocols.

## Author contributions

S.J.V. and J.F.J. developed the concepts, conceived of the study plan, and wrote the manuscript.

S.J.V. designed and performed the experiments with assistance from S.A.P., V.A.S.E., A.GM., G.X.C., A.R., A.C.L., E.S.N., P.K., J.D.F., K.G., and D.T.S. J.F.J provided project administration, supervision, resources, and data analysis, the latter with V.A.S.E. and S.J.V.’s assistance. A.B. performed 3D structure modeling and analysis of protein interactions under the supervision of F.X.V., who wrote the corresponding section of the manuscript. A.GM. perform molecular docking simulations.

