## Supplementary material for "Expanding salivary biomarker detection by creating a synthetic neuraminic acid sensor via chimeragenesis": Suppl_Materials

### INDEX

#### Supplementary Tables

- ***Supplementary Table 1.*** Linker (LNK) regions used in this study.
- ***Supplementary Table 2.*** Bacterials strains and plasmids used in this study.
- ***Supplementary Table 3.*** Oligonucleotides used in this study.
- ***Supplementary Table 4.*** Relative fluorescence over time of different MG1655 (pCKT-*Chimera*, pHC_DYOLacI-R) strains.

#### Supplementary Figures

- ***Supplementary Figure 1.*** Growth of MG1655 (pHC_DYOLacI-R) supplemented with different concentrations of neuraminic acid.
- ***Supplementary Figure 2.*** *In vivo* behaviour of MG1655 (pCKT-Siren, pHC_DYOLacI-R) strain expressing Siren (LacI-LNK1-SiaP).
- ***Supplementary Figure 3.*** *In vivo* behaviour of MG1655 (pCKT-Kunst, pHC_DYOLacI-,) strain expressing Kunst (LacI-LNK3-SiaP).
- ***Supplementary Figure 4***. qRT-PCR analysis of the regulated expression of GFP by Sphnx.

**Appendix:** Relevant transcription factor sequences.

### Supplementary Table 1

***Supplementary Table 1.*** Linker (LNK) regions used in this study.

| Linker | Encoding sequence | Amino acid sequence | Alternative names | Reference or Source |
|---|---|---|---|---|
| LNK1 | – | – | No linker | - |
| LNK2 | GAGAAGGAGAAAGAGAAG | EKEKEK | Ben_1 | 1 |
| LNK3 | GGTAGCGGCAGCGGTAGC | (GS)3 | BBa_J18921 | 2 |
| LNK4 | GGTGGAGGAGGTTCTGGAGGCGGTGGAAGTGGTGGCGGAGGTAGC | (GGGGS)5 | BBa_K157013 | 2 |
| LNK5 | GGTGGTTCTGGT | GGSG | BBa_K243004 | 2 |
| LNK6 | TCGTTGCTGATTGGCGTT | SLLIGV | DBD-LacI 61-66 | - |

**Table References**


1 Juárez *et al.* (2018). Biosensor libraries harness large classes of binding domains for construction of allosteric transcriptional regulators. Nat Commun. 6;9(1):3101[PMID 30082754].

2 iGEM Registry of Standard Biological Parts (Registry): parts.igem.org (Access June 8, 2024)

### Supplementary Table 2

***Supplementary Table 2.*** Bacterials strains and plasmids used in this study.

| Strain or Plasmid | Description | Ref. or Source |
|---|---|---|
| ***Escherichia coli* strains** | | |
| NEB5-alpha | *huA2* $\Delta$*(argF-lacZ)U169 phoA glnV44* $\Phi 80\Delta$*(lacZ)M15 gyrA96 recA1 relA1 endA1 thi-1 hsdR17* | NEB, Ipswich, MA |
| MG1655 | $F^- \lambda^-$ *ilvG- rfb-50 rph-1* | 1 |
| MC4100 | $F^- \lambda^-$ *araD139* $\Delta$*(argF-lac)U169 rpsL150* $(Sm^R)$ *relA1 flbB5301 deoC1 ptsF25 rbsR* | 2 |
| **Plasmids** | | |
| **pUC19** | $Ap^R$; *oriColE1* , cloning vector | 3 |
| pHC_DYOLacI-R | $Ap^R$; Reporter plasmid containing $P_{lac}$ *-sfGFP* for the screening of TFs carrying DBD-LacI. | 4 |
| pUC19RBS | pUC19 derivative includinga consensus RBS downstream of $P_{lac}$ . | This work |
| pUC19-Chimera | Family of vectors expressing chimeric transcription factors | |
| pUC19-Siren | $Ap^R$; pUC19 derivative expressing LacI-LNK1-SiaP (no linker) chimera, also known as Siren | This work |
| pUC19-Sphnx | $Ap^R$; pUC19 derivative expressing LacI-LNK2-SiaP chimera, also known as Sphnx | This work |
| pUC19-Kunst | $Ap^R$; pUC19 derivative expressing LacI-LNK3-SiaP chimera, also known as Kunst | This work |
| pUC19-Bastet | $Ap^R$; pUC19 derivative expressing LacI-LNK5-SiaP chimera, also known as Bastet | This work |
| pUC19-Lumos | $Ap^R$; pUC19 derivative expressing LacI-LNK6-SiaP (duplication) chimera, also known as Lumos | This work |
| pUC19-Harpy | $Ap^R$; pUC19 derivative expressing LacI-LNK1-SatA chimera, also known as Harpy | This work |
| pUC19-Haus | $Ap^R$; pUC19 derivative expressing LacI-LNK2-SatA chimera, also known as Haus | This work |
| **pCKTRBS** | $Cm^R$; pCK01 (*oripSC101*) derivative carrying *tetR* under a constitutive promoter and an aTc-inducible $P_{tetO}$ promoter for control expression of genes. | 4 |
| pCKTRBS-LacIwt | $Cm^R$; pCKTRBS carrying a wild-type LacI | 4 |
| pCKT-Chimera | Family of vectors expressing chimeric transcription factors | |
| pCKT-Siren | $Cm^R$; pCKTRBS derivative expressing LacI-LNK1-SiaP (no linker) chimera, also known as Siren | This work |
| pCKT-Sphnx | $Cm^R$; pCKTRBS derivative expressing LacI-LNK2-SiaP chimera, also known as Sphnx | This work |
| pCKT-Kunst | $Cm^R$; pCKTRBS derivative expressing LacI-LNK3-SiaP chimera, also known as Kunst | This work |
| pCKT-Bastet | $Cm^R$; pCKTRBS derivative expressing LacI-LNK5-SiaP chimera, also known as Bastet | This work |
| pCKT-Lumos | $Cm^R$; pCKTRBS derivative expressing LacI-LNK6-SiaP (duplication) chimera, also known as Lumos | This work |
| pCKT-Harpy | $Cm^R$; pCKTRBS derivative expressing LacI-LNK1-SatA chimera, also known as Harpy | This work |
| pCKT-Haus | $Cm^R$; pCKTRBS derivative expressing LacI-LNK2-SatA chimera, also known as Haus | This work |

**Table References**


1 Blattner *et al.* (1997). The complete genome sequence of *Escherichia coli* K-12. Science. 277(5331):1453-62 [PMID 9278503].

2 Casadaban (1976). Transposition and fusion of the *lac* genes to selected promoters in *Escherichia coli* using bacteriophage *lambda* and *Mu* . J Mol Biol. 104(3):541-55 [PMID 781293].

3 Yanisch-Perron *et al.* (1985) Improved M13 phage cloning vectors and host strains: nucleotide sequences of the M13mp18 and pUC19 vectors. Gene 33, 103–119.

4 Juárez *et al.* (2018). Biosensor libraries harness large classes of binding domains for construction of allosteric transcriptional regulators. Nat Commun. 6;9(1):3101[PMID 30082754].

### Supplementary Table 3

***Supplementary Table 3.*** Oligonucleotides used in this study.

| Oligonucleotide name | Sequence | Purpose / Comments |
|---|---|---|
| LacI_RBS_Tail_5_SV00001 | CGGTACCCGGGTGACCTAAGGAGGTAAATAATGAAACCAGTAACGTTATACGATGTCG | Amplification of *lacI* DNA binding domain (*DBD-lacI*). |
| LacI_3_SV00002 | AACGCCAATCAGCAACGACTGTTTGCCCG | |
| satA_LNK1_F1_SV00030 | GCGGGCAAACAGTCGTTGCTGATTGGCGTTATGGCTGCCAATCTGCATATTTTCGAAG | Amplification of *satA* (LNK1-LBD) without including any linker sequence in the 5' overhang. |
| satA_LNK2_F1_SV00031 | GCGGGCAAACAGTCGTTGCTGATTGGCGTTGAGAAGGAGAAAGAGAAGATGGCTGCCAATCTGCATATTTTCGAAG | Amplification of *satA* (LNK2-LBD) including LNK2 encoding sequence in the 5' overhang. |
| satA_LNK3_F1_SV00032 | GCGGGCAAACAGTCGTTGCTGATTGGCGTTGGTAGCGGCAGCGGTAGCATGGCTGCCAATCTGCATATTTTCGAAG | Amplification of *satA* (LNK3-LBD) including LNK3 encoding sequence in the 5' overhang. |
| satA_LNK4_F1_SV00033 | GCGGGCAAACAGTCGTTGCTGATTGGCGTTGGAGGCGGTGGAAGTGGTGGCGGAGGTAGCATGGCTGCCAATCTGCATATTTTCGAAG | Amplification of *satA* (LNK4-LBD) including LNK4 encoding sequence in the 5' overhang. |
| satA_LNK5_F1_SV00034 | GCGGGCAAACAGTCGTTGCTGATTGGCGTTGGTGGTTCTGGTATGGCTGCCAATCTGCATATTTTCGAAG | Amplification of *satA* (LNK5-LBD) including LNK5 encoding sequence in the 5' overhang. |
| satA_END_3_SV00004 | TGCATGCCTGCAGGTCCACTCTAGAGGATCTTATGTACGACCTACACCAAGGAAAGACAA | Amplification of every *satA* variant. |
| siaP_LNK1_5_SV00015 | GCGGGCAAACAGTCGTTGCTGATTGGCGTTATGATGAAATTGACAAAACTTTTCCTTGCC | Amplification of *siaP* (LNK1-LBD) without including any linker sequence in the 5' overhang. |
| siaP_LNK2_5_SV00016 | GCGGGCAAACAGTCGTTGCTGATTGGCGTTGAGAAGGAGAAAGAGAAGATGATGAAATTGACAAAACTTTTCCTTGCC | Amplification of *siaP* (LNK2-LBD) including LNK2 encoding sequence in the 5' overhang. |
| siaP_LNK3_5_SV00017 | GCGGGCAAACAGTCGTTGCTGATTGGCGTTGGTAGCGGCAGCGGTAGCATGATGAAATTGACAAAACTTTTCCTTGCC | Amplification of *siaP* (LNK3-LBD) including LNK3 encoding sequence in the 5' overhang. |
| siaP_LNK4_5_SV00018 | GCGGGCAAACAGTCGTTGCTGATTGGCGTTGGAGGCGGTGGAAGTGGTGGCGGAGGTAGCATGATGAAATTGACAAAACTTTTCCTTGCC | Amplification of *siaP* (LNK4-LBD) including LNK4 encoding sequence in the 5' overhang. |
| siaP_LNK5_5_SV00019 | GCGGGCAAACAGTCGTTGCTGATTGGCGTTTCGTTGCTGATTGGCGTTATGATGAAATTG | Amplification of *siaP* (LNK5-LBD) including LNK5 encoding sequence in the 5' overhang. |
| siaP_END_3_v2_JFJ0018 | CCTGCAGGTCCACTCTAGAGGATCTTATGGATTGATTGCTTCAATTTGTTTTAAAGCTGATTCACC | Amplification of every *siaP* variant. |
| pUC_3'_RBS_Tail_JFJ0019 | TATTTACCTCCTTAGGTCACCCGGGTACCGGCCTGGGGTGCCTAATGAGTGAGCTAACTC | Amplification of pUC19 (divergent PCR) introducing a consensus RBS sequence to originate pUC19RBS fragment. |
| pUC_5'_END_Tail_JFJ0020 | GATCCTCTAGAGTGGACCTGCAGGCATGCATTAAGCCAGCCCCGACACCCG | |
| pCKTRBS 3'.LacI_SV00022 | TGCGACATCGTATAACGTTACTGGTTTCATTATTTACCTCCTTAGGTCACCC | Amplification of pCKTRBS (divergent PCR) introducing homology arms to *lacI* and *satA* / *siaP*. |
| pCKTRBS 5'.SatA_SV00023 | CTTGGTGTAGGTCGTACATAAGATCCTCTAGAGTGGACCTGCAGG | |
| pCKTRBS 5'.SiaP_SV00021 | TTAAAACAAATTGAAGCAATCAATCCATAAGATCCTCTAGAGTGGACCTGCAGGCATGCA | |
| LacI_F1_SV00024 | ATGAAACCAGTAACGTTATACGATGTCGCA | Amplification of assembled chimeric genes. |
| SiaP_R1_SV00025 | TTATGGATTGATTGCTTCAATTTGTTTTTGACCAGT | |
| SatA_R1_SV00026 | TTATGTACGACCTACACCAAGGAAAGACAA | |
| qPCR_Consensus_RBS-sfGFP_5'_SV00027A | AGGTAAATAATGCGTAAAGGCGAAGAG | qRT-PCR amplification of a transcribed region encompasing the RBS and the beginning of *sfgfp* in pHC_DYOLacI-R |
| qPCR_sfGFP_3'_SV00027 | AGTCATGCTGCTTCATGTGGTC | |
| gel_shift_pHC_DYOLacI_Upstream_Ptac_5'_SV00028 | CTGAAATGAGCTGTTGACAATTAATCATC | Amplification of a dsDNA probe for gel shift (EMSA) spanning the $P_{tac}$ promoter of pHC_DYOLacI-R including *lacO*. |
| gel_shift_sfGFP_3'.2_SV00029 | AGTAGTACAGATGAACTTCAGCGTC | |

### Supplementary Table 4

| Name | | |
|---|---|---|
| **Common** | **Systematic** | **Graphs** |
| **Siren** | LacI-LNK1-SiaP | 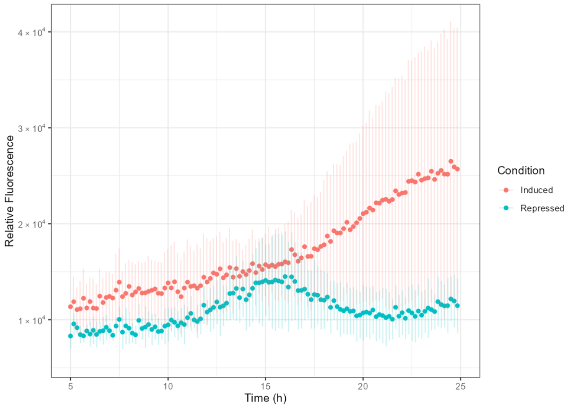  |
| **Sphnx** | LacI-LNK2-SiaP | 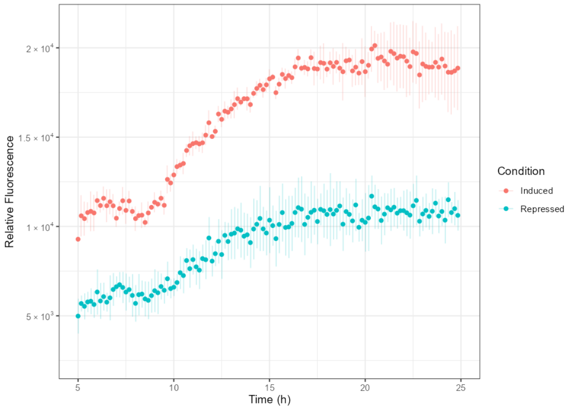  |
| **Kunst** | LacI-LNK3-SiaP | 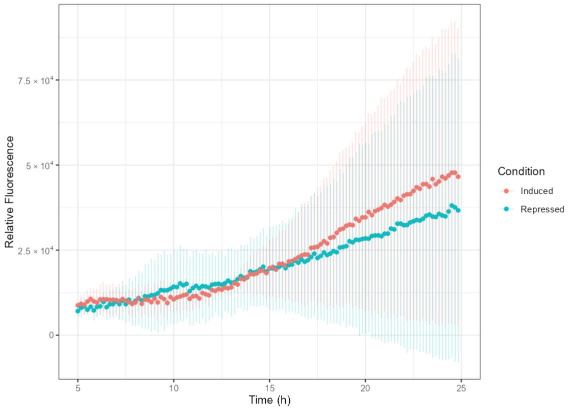  |

<table>
<tr><td>Scorpio</td><td>LacI-LNK4-SiaP</td><td>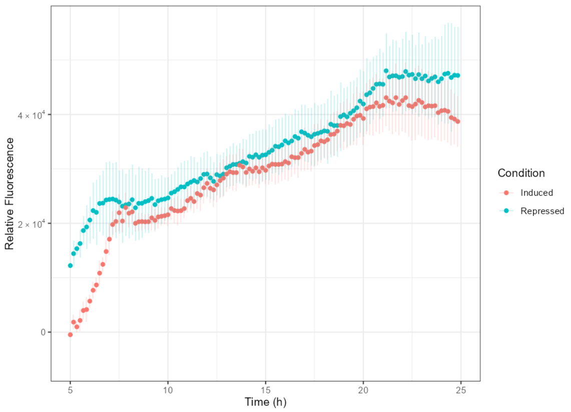
</td></tr>
<tr><td>Bastet</td><td>LacI-LNK5-SiaP</td><td>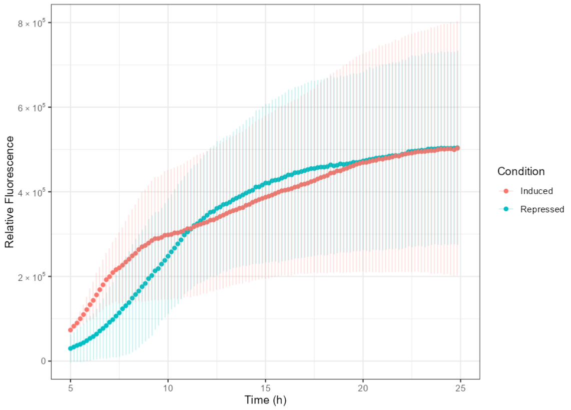
</td></tr>
<tr><td>Lumos</td><td>LacI-LNK6-SiaP</td><td>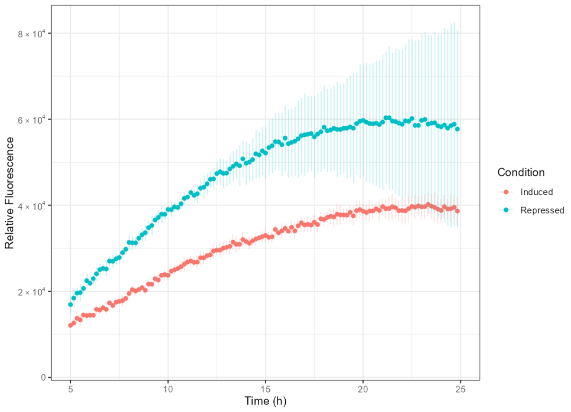
</td></tr>
</table>

| | | |
|---|---|---|
| **Harpy** | LacI-LNK1-SatA | 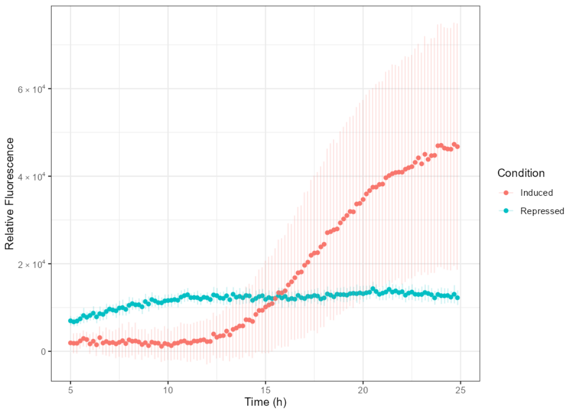 |
| **Haus** | LacI-LNK2-SatA | 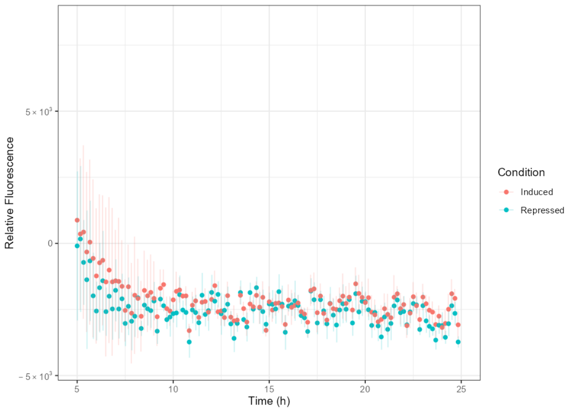 |

***Supplementary Table 4.*** Relative fluorescence over time of different MG1655 (pCKT-*Chimera*, pHC_DYOLacI-R) strains. The first two columns of this summary table includes both the systematic and common names assigned to every chimeric TF created in this work. The third column displays a representative time course (hours) showing relative fluorescence of MG1655 (pCKT-*Chimera*, pHC_DYOLacI-R) cells grown in a multiwell plate reader, with error bars denoting SEM (standard error of the mean) associated to technical replicas (*n* = 4). Promoter activity is measured as relative fluorescence (GFP-associated fluorescence in arbitrary units / $OD_{600}$) of the strain growing in the de-repressed (aTc$^+$, Neu5Ac$^+$; *orange dots*) or repressed (aTc$^+$, Neu5Ac$^-$; *blue dots*) states.

### Supplementary Figure 1

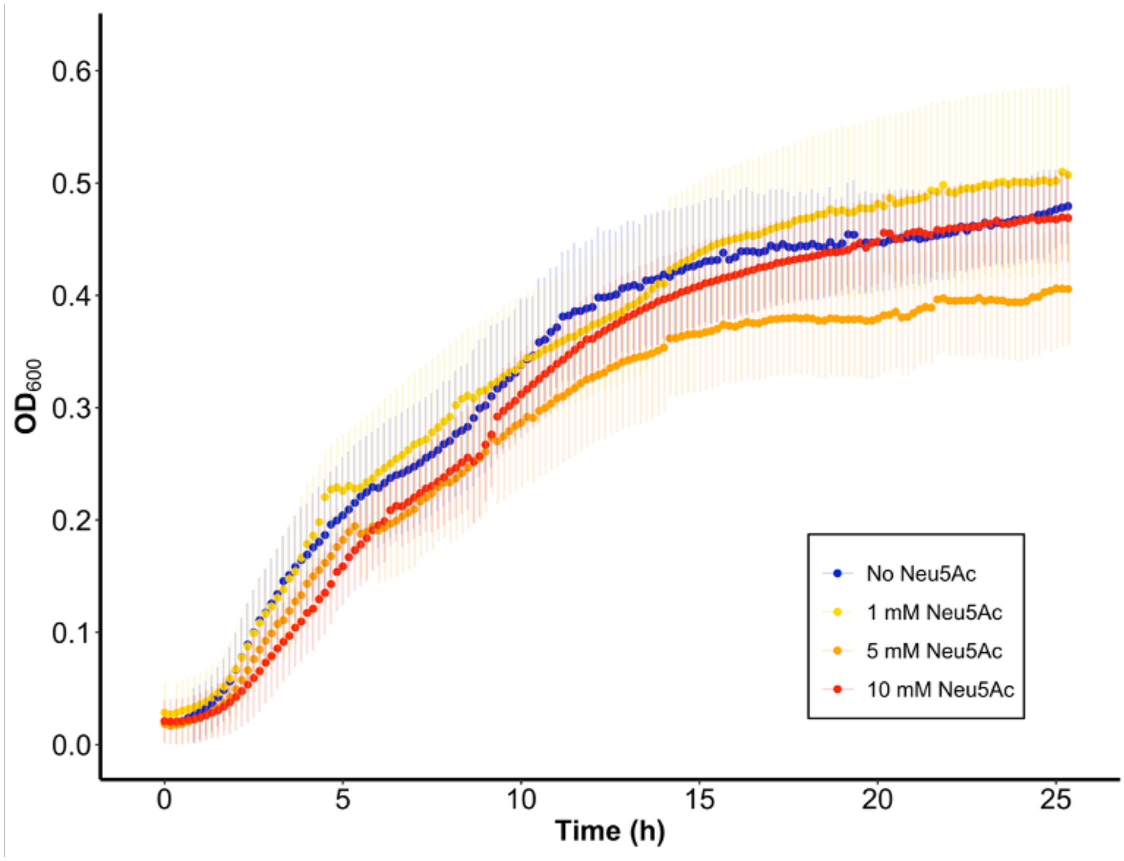


***Supplementary Figure 1.*** Growth of MG1655 (pHC_DYOLacI-R) in LB media (*green dots*) or in LB media supplemented with neuraminic acid at concentrations: 1 mM (*yellow dots*), 5 mM (*orange dots*), 10 mM (*red dots*). Bacterial cultures were grown and their $OD_{600}$ and GFP-associated fluorescence tracked over time in a multiwell plate reader as described in *Materials and Methods.* Error bars denote SEM (standard error of the mean) associated to independent biological replicas ($n$ = 3).

### Supplementary Figure 2

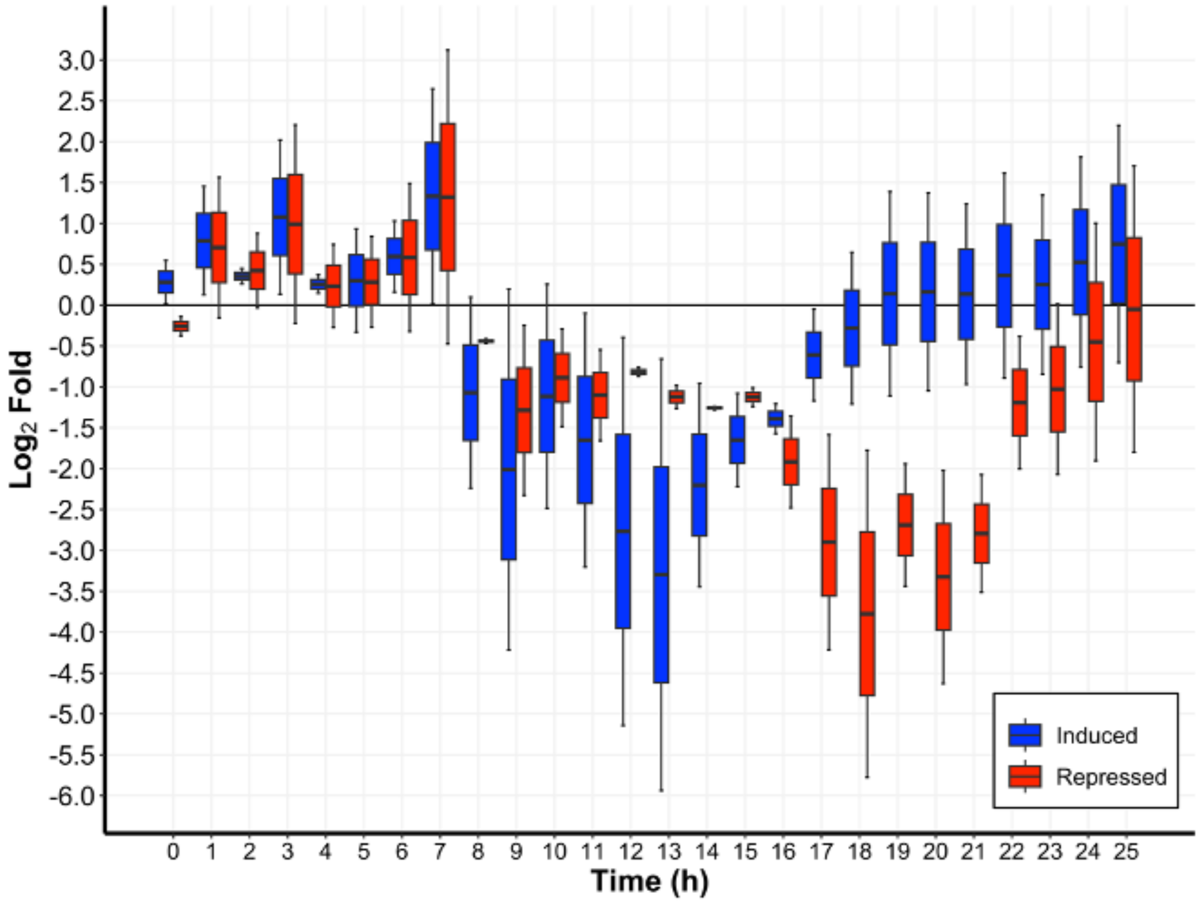


***Supplementary Figure 2. In vivo* behaviour of a strain expressing Siren (LacI-LNK1-SiaP).** Time course (hours) showing relative fluorescence of MG1655 (pCKT-Siren, pHC_DYOLacI-R) cells grown in a multiwell plate reader. Promoter activity is measured as $log_2$ fold of the relative fluorescence (GFP-associated fluorescence in arbitrary units / $OD_{600}$) of the strain growing in the de-repressed ($aTc^+$, $Neu5Ac^+$; *blue boxplots*) or repressed ($aTc^+$, $Neu5Ac^-$; *red boxplots*) states compared to the basal expression of the reporter ($aTc^-$, $Neu5Ac^-$). Boxplots with whiskers represent data dispersion of the average values of biological replicas (*n* = 8). It can be appreciated how in order to approach the basal activity ($log_2$ fold = 0) around 17 - 21 h, the addition of neuraminic acid is necessary when the chimera gets expressed by the supplementation of aTc. Growth conditions and fluorescence assays performed as described in *Materials and Methods*.

### Supplementary Figure 3.

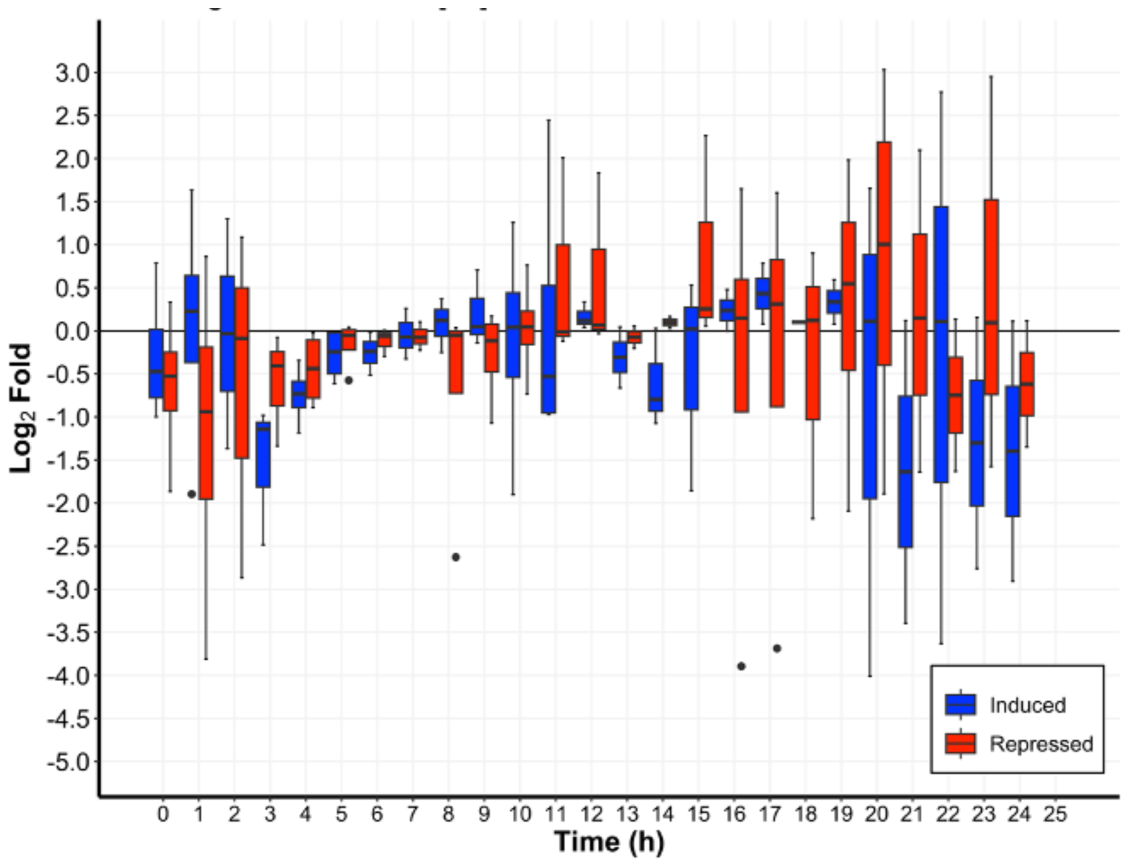


***Supplementary Figure 3*. *In vivo* behaviour of a strain expressing Kunst (LacI-LNK3-SiaP).** Time course (hours) showing relative fluorescence of MG1655 (pCKT-Kunst, pHC_DYOLacI-R) cells grown in a multiwell plate reader. Promoter activity is measured as $\log_2$ fold of the relative fluorescence (GFP-associated fluorescence in arbitrary units / $OD_{600}$) of the strain growing in the de-repressed ($aTc^+$, $Neu5Ac^+$; *blue boxplots*) or repressed ($aTc^+$, $Neu5Ac^-$; *red boxplots*) states compared to the basal expression of the reporter ($aTc^-$, $Neu5Ac^-$). Boxplots with whiskers represent data dispersion of the average values of biological replicas ($n$ = 8). It can be appreciated how the strain is not able to display GFP-associated fluorescence in the presence of neuraminic acid. Growth conditions and fluorescence assays performed as described in *Materials and Methods*.

### Supplementary Figure 4

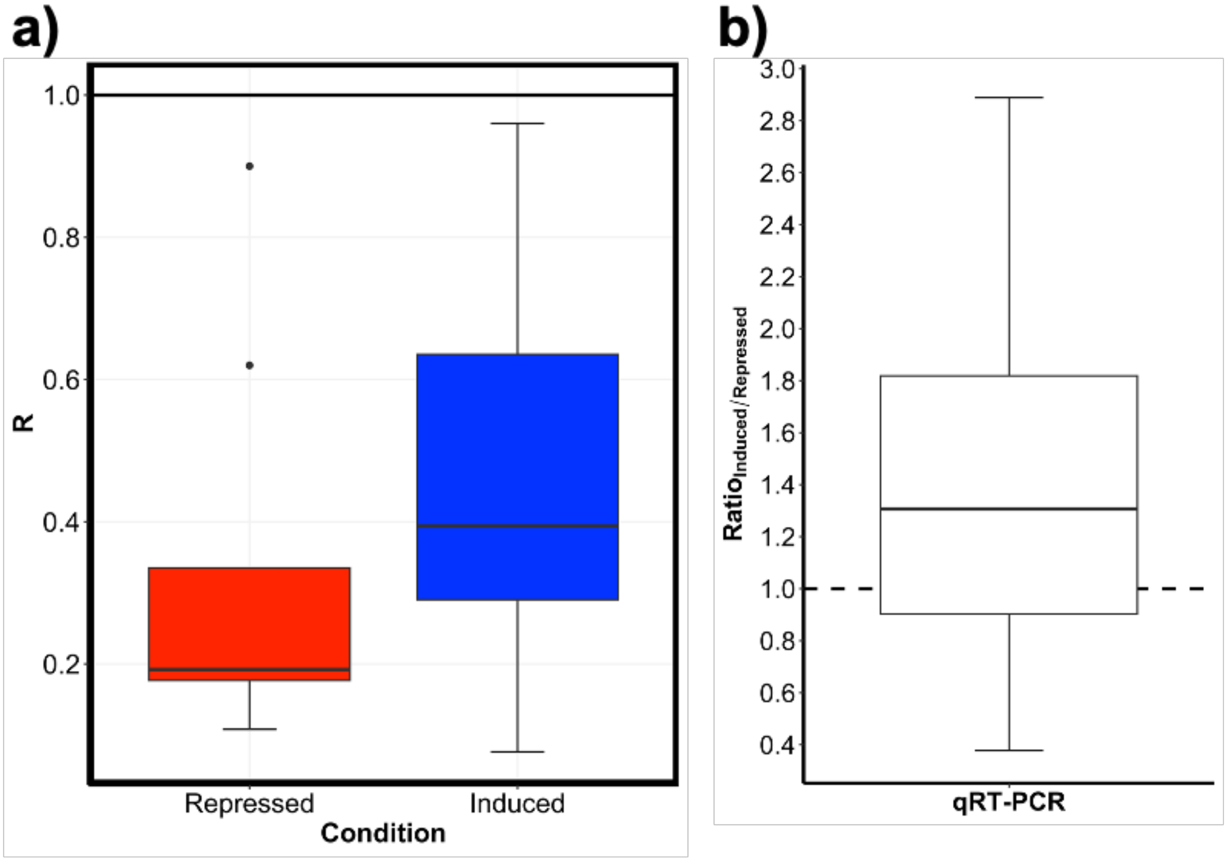


***Supplementary Figure 4.*** Relative activity of the $P_{lac}$ promoter, regulated by Sphnx, measured by quantitative real-time PCR (qRT-PCR). Total RNA was isolated from MG1655 (pCKT-Sphnx, pHCDYOLacI-R) cultures performed as detailed in *Materials and Methods*. **a)** Boxplot depicting the relative activity of the $P_{lac}$ promoter expressed as fold gene expression ($R = 2^{-\Delta\Delta Ct}$) in cells grown under repression (aTc$^{+}$/Neu5Ac$^{-}$; *red*) or de-repression (aTc$^{+}$/Neu5Ac$^{+}$; *blue*) conditions using the basal state (aTc$^{-}$/Neu5Ac$^{-}$) as a reference ($n$ = 8). **b)** Graph showing the induced to repressed ratio ($R_{induced}$ / $R_{repressed}$) for the data displayed in panel a).

### Appendix: Relevant transcription factor sequences

Common name: **Sphnx**.

Systematic name: **LacI**-**LNK2**-**SiaP** ($N_t$-DBD-LNK-LBD-$C_t$)

```
>sphnx_nt_sequence (1206 bp)
ATGAAACCAGTAACGTTATACGATGTCGCAGAGTATGCCGGTGTCTCTTATCAGACCGTTTCCC
GCGTGGTGAACCAGGCCAGCCACGTTTCTGCGAAAACGCGGGAAAAAGTGGAAGCGGCGATGGC
GGAGCTGAATTACATTCCCAACCGCGTGGCACAACAACTGGCGGGCAAACAGTCGTTGCTGATT
GGCGTTGAGAAGGAGAAAGAGAAGATGATGAAATTGACAAAACTTTTCCTTGCCACCGCCATTT
CTTTAGGCGTATCTTCTGCTGTTCTTGCCGCTGATTATGACTTGAAATTCGGTATGAATGCTGG
AACTTCATCAAATGAATATAAAGCGGCAGAAATGTTTGCCAAAGAAGTCAAAGAAAAATCACAG
GGTAAAATTGAAATTTCACTTTATCCAAGTTCACAATTAGGTGATGACCGTGCAATGTTAAAAC
AATTAAAAGACGGTTCTCTCGACTTTACCTTTGCAGAATCTGCTCGCTTCCAGCTGTTTTACCC
TGAAGCGGCAGTATTTGCCTTACCTTATGTTATTAGCAACTACAATGTTGCACAAAAAGCCTTA
TTCGATACAGAATTCGGTAAAGATTTAATTAAAAAAATGGATAAAGATCTTGGCGTGACTTTAC
TTTCCCAAGCTTATAACGGAACTCGCCAAACGACTTCAAATCGTGCAATCAACAGTATTGCAGA
TATGAAAGGCTTAAAACTTCGTGTGCCAAATGCAGCAACAAACTTAGCCTATGCTAAATATGTT
GGTGCATCACCAACACCAATGGCATTTTCTGAAGTTTATCTTGCGTTACAAACCAATGCCGTCG
ATGGTCAAGAAAACCCGTTAGCAGCGGTGCAAGCACAAAAATTCTATGAAGTGCAAAAGTTCTT
AGCAATGACTAATCATATTTTGAATGACCAACTTTATTTAGTAAGCAACGAGACTTATAAAGAA
CTCCCTGAAGATCTTCAAAAAGTCGTAAAAGATGCTGCCGAAAATGCAGCAAAATATCACACTA
AATTATTCGTAGATGGAGAGAAAGATTTAGTCACATTCTTTGAAAAACAAGGCGTGAAAATTAC
ACATCCTGATCTTGTTCCATTTAAAGAATCAATGAAGCCGTATTATGCTGAGTTTGTAAAACAA
ACTGGTCAAAAAGGTGAATCAGCTTTAAAACAAATTGAAGCAATCAATCCATAA
>Sphnx_aa_sequence (401 aa)
MKPVTLYDVAEYAGVSYQTVSRVVNQASHVSAKTREKVEAAMAELNYIPNRVAQQLAGKQSLLI
GVEKEKEKMMKLTKLFLATAISLGVSSAVLAADYDLKFGMNAGTSSNEYKAAEMFAKEVKEKSQ
GKIEISLYPSSQLGDDRAMLKQLKDGSLDFTFAESARFQLFYPEAAVFALPYVISNYNVAQKAL
FDTEFGKDLIKKMDKDLGVTLLSQAYNGTRQTTSNRAINSIADMKGLKLRVPNAATNLAYAKYV
GASPTPMAFSEVYLALQTNAVDGQENPLAAVQAQKFYEVQKFLAMTNHILNDQLYLVSNETYKE
LPEDLQKVVKDAAENAAKYHTKLFVDGEKDLVTFFEKQGVKITHPDLVPFKESMKPYYAEFVKQ
TGQKGESALKQIEAINP
```

Common name: **Kunst**.

Systematic name: **LacI-LNK3-SiaP** ($N_t$-DBD-LNK-LBD-$C_t$)

```
>kunst_nt_sequence (1206 bp)
ATGAAACCAGTAACGTTATACGATGTCGCAGAGTATGCCGGTGTCTCTTATCAGACCGTTTCCC
GCGTGGTGAACCAGGCCAGCCACGTTTCTGCGAAAACGCGGGAAAAAGTGGAAGCGGCGATGGC
GGAGCTGAATTACATTCCCAACCGCGTGGCACAACAACTGGCGGGCAAACAGTCGTTGCTGATT
GGCGTTGGTAGCGGCAGCGGTAGCATGATGAAATTGACAAAACTTTTCCTTGCCACCGCCATTT
CTTTAGGCGTATCTTCTGCTGTTCTTGCCGCTGATTATGACTTGAAATTCGGTATGAATGCTGG
AACTTCATCAAATGAATATAAAGCGGCAGAAATGTTTGCCAAAGAAGTCAAAGAAAAATCACAG
GGTAAAATTGAAATTTCACTTTATCCAAGTTCACAATTAGGTGATGACCGTGCAATGTTAAAAC
AATTAAAAGACGGTTCTCTCGACTTTACCTTTGCAGAATCTGCTCGCTTCCAGCTGTTTTACCC
TGAAGCGGCAGTATTTGCCTTACCTTATGTTATTAGCAACTACAATGTTGCACAAAAAGCCTTA
TTCGATACAGAATTCGGTAAAGATTTAATTAAAAAAATGGATAAAGATCTTGGCGTGACTTTAC
TTTCCCAAGCTTATAACGGAACTCGCCAAACGACTTCAAATCGTGCAATCAACAGTATTGCAGA
TATGAAAGGCTTAAAACTTCGTGTGCCAAATGCAGCAACAAACTTAGCCTATGCTAAATATGTT
GGTGCATCACCAACACCAATGGCATTTTCTGAAGTTTATCTTGCGTTACAAACCAATGCCGTCG
ATGGTCAAGAAAACCCGTTAGCAGCGGTGCAAGCACAAAAATTCTATGAAGTGCAAAAGTTCTT
AGCAATGACTAATCATATTTTGAATGACCAACTTTATTTAGTAAGCAACGAGACTTATAAAGAA
CTCCCTGAAGATCTTCAAAAAGTCGTAAAAGATGCTGCCGAAAATGCAGCAAAATATCACACTA
AATTATTCGTAGATGGAGAGAAAGATTTAGTCACATTCTTTGAAAAACAAGGCGTGAAAATTAC
ACATCCTGATCTTGTTCCATTTAAAGAATCAATGAAGCCGTATTATGCTGAGTTTGTAAAACAA
ACTGGTCAAAAAGGTGAATCAGCTTTAAAACAAATTGAAGCAATCAATCCATAA
>Kunst_aa_sequence (401 aa)
MKPVTLYDVAEYAGVSYQTVSRVVNQASHVSAKTREKVEAAMAELNYIPNRVAQQLAGKQSLLI
GVGSGSGSMMKLTKLFLATAISLGVSSAVLAADYDLKFGMNAGTSSNEYKAAEMFAKEVKEKSQ
GKIEISLYPSSQLGDDRAMLKQLKDGSLDFTFAESARFQLFYPEAAVFALPYVISNYNVAQKAL
FDTEFGKDLIKKMDKDLGVTLLSQAYNGTRQTTSNRAINSIADMKGLKLRVPNAATNLAYAKYV
GASPTPMAFSEVYLALQTNAVDGQENPLAAVQAQKFYEVQKFLAMTNHILNDQLYLVSNETYKE
LPEDLQKVVKDAAENAAKYHTKLFVDGEKDLVTFFEKQGVKITHPDLVPFKESMKPYYAEFVKQ
TGQKGESALKQIEAINP
```

Common name: **Siren**.

Systematic name: **LacI-LNK1-SiaP** ($N_t$-DBD-LBD-$C_t$), no linker sequence

>*siren*_nt_sequence (1188 bp)

ATGAAACCAGTAACGTTATACGATGTCGCAGAGTATGCCGGTGTCTCTTATCAGACCGTTTCCC
GCGTGGTGAACCAGGCCAGCCACGTTTCTGCGAAAACGCGGGAAAAAGTGGAAGCGGCGATGGC
GGAGCTGAATTACATTCCCAACCGCGTGGCACAACAACTGGCGGGCAAACAGTCGTTGCTGATT
GGCGTTATGATGAAATTGACAAAACTTTTCCTTGCCACCGCCATTTCTTTAGGCGTATCTTCTG
CTGTTCTTGCCGCTGATTATGACTTGAAATTCGGTATGAATGCTGGAACTTCATCAAATGAATA
TAAAGCGGCAGAAATGTTTGCCAAAGAAGTCAAAGAAAAATCACAGGGTAAAATTGAAATTTCA
CTTTATCCAAGTTCACAATTAGGTGATGACCGTGCAATGTTAAAACAATTAAAAGACGGTTCTC
TCGACTTTACCTTTGCAGAATCTGCTCGCTTCCAGCTGTTTTACCCTGAAGCGGCAGTATTTGC
CTTACCTTATGTTATTAGCAACTACAATGTTGCACAAAAAGCCTTATTCGATACAGAATTCGGT
AAAGATTTAATTAAAAAAATGGATAAAGATCTTGGCGTGACTTTACTTTCCCAAGCTTATAACG
GAACTCGCCAAACGACTTCAAATCGTGCAATCAACAGTATTGCAGATATGAAAGGCTTAAAACT
TCGTGTGCCAAATGCAGCAACAAACTTAGCCTATGCTAAATATGTTGGTGCATCACCAACACCA
ATGGCATTTTCTGAAGTTTATCTTGCGTTACAAACCAATGCCGTCGATGGTCAAGAAAACCCGT
TAGCAGCGGTGCAAGCACAAAAATTCTATGAAGTGCAAAAGTTCTTAGCAATGACTAATCATAT
TTTGAATGACCAACTTTATTTAGTAAGCAACGAGACTTATAAAGAACTCCCTGAAGATCTTCAA
AAAGTCGTAAAAGATGCTGCCGAAAATGCAGCAAAATATCACACTAAATTATTCGTAGATGGAG
AGAAAGATTTAGTCACATTCTTTGAAAAACAAGGCGTGAAAATTACACATCCTGATCTTGTTCC
ATTTAAAGAATCAATGAAGCCGTATTATGCTGAGTTTGTAAAACAAACTGGTCAAAAAGGTGAA
TCAGCTTTAAAACAAATTGAAGCAATCAATCCATAA

>Siren_aa_sequence (396 aa)

MKPVTLYDVAEYAGVSYQTVSRVVNQASHVSAKTREKVEAAMAELNYIPNRVAQQLAGKQSLLI
GVMMKLTKLFLATAISLGVSSAVLAADYDLKFGMNAGTSSNEYKAAEMFAKEVKEKSQGKIEIS
LYPSSQLGDDRAMLKQLKDGSLDFTFAESARFQLFYPEAAVFALPYVISNYNVAQKALFDTEFG
KDLIKKMDKDLGVTLLSQAYNGTRQTTSNRAINSIADMKGLKLRVPNAATNLAYAKYVGASPTP
MAFSEVYLALQTNAVDGQENPLAAVQAQKFYEVQKFLAMTNHILNDQLYLVSNETYKELPEDLQ
KVVKDAAENAAKYHTKLFVDGEKDLVTFFEKQGVKITHPDLVPFKESMKPYYAEFVKQTGQKGE
SALKQIEAINP
